# Ongoing production of low-fitness hybrids limits range overlap between divergent cryptic species

**DOI:** 10.1101/2021.02.09.430089

**Authors:** Else K. Mikkelsen, Darren Irwin

## Abstract

Contact zones between recently-diverged taxa provide opportunities to examine the causes of reproductive isolation and to examine the processes that determine whether two species can coexist over a broad region. The Pacific Wren (*Troglodytes pacificus*) and the Winter Wren *(Troglodytes hiemalis*) are two morphologically similar songbird species that started diverging about 4 million years ago, older than most sister species pairs. The ranges of these species come into narrow contact in western Canada, where the two species remain distinct in sympatry. To assess evidence for differentiation, hybridization, and introgression in this system, we examined variation in over 250,000 single nucleotide polymorphism markers distributed across the genomes of the two species. The two species formed highly divergent genetic clusters, consistent with long-term differentiation. In a set of 75 individuals from allopatry and sympatry, two first-generation hybrids (i.e., F1’s) were detected, indicating only moderate levels of assortative mating between these taxa. We found no recent backcrosses or F2’s or other evidence of recent breeding success of F1 hybrids, indicating very low or zero fitness of F1 hybrids. Examination of genomic variation shows evidence for only a single backcrossing event in the distant past. The sizeable rate of hybridization combined with very low fitness of F1 hybrids is expected to result in a population sink in the contact zone, largely explaining the narrow overlap of the two species. If such dynamics are common in nature, they could explain the narrow range overlap often observed between pairs of closely related species. Additionally, we present evidence for a rare duplication of a large chromosomal segment from an autosome to the W chromosome, the female-specific sex chromosome in birds.

## Introduction

One of the foundational goals of evolutionary biology is to understand the mechanisms at play as one species splits into two. A major debate in the speciation literature (e.g., Mayr, 1942; Price, 2008) has centred on the relative importance of assortative mating (i.e., premating isolation) and low hybrid fitness (i.e., postzygotic isolation). An often-articulated view is that premating isolation is most important during early stages of the speciation process, with postzygotic isolation building later (Mayr, 1942; Grant & Grant, 1997; Schumer et al., 2017). Also debated is how important geographic separation of populations (i.e., allopatry) is in causing speciation, versus the possibility of sympatric speciation or speciation-with-gene-flow (e.g., Nosil, 2008). An often-observed pattern of closely related species occurring in neighbouring geographic regions with just a small region of overlap implies that some process prevents coexistence between species. This restricted overlap is often interpreted to be a result of differential specialization to distinct habitats that change across space, and/or niche exclusion, in which species are so similar they compete for the same resources (Price & Kirkpatrick, 2009; Lee-Yaw & Irwin, 2015). Hybridization and/or reproductive interference can also limit range overlap between closely related species (Barton & Hewitt, 1989; Gröning & Hochkirch, 2008). Close examination of the overlap zone between closely related species can provide insight into the strength and causes of reproductive isolation as well as the reasons for limited range overlap.

Recent advancements in DNA sequencing technology (Elshire et al., 2011; Goodwin, McPherson, & McCombie, 2016; Heather & Chain, 2016) have enabled the detection of differentiation, hybridization, and genetic introgression between diverging species (Payseur & Rieseberg, 2016). Genotyping-by-sequencing (GBS; Elshire et al. 2011) allows for many individuals to be simultaneously genotyped at tens or hundreds of thousands of genetic loci, providing genome-wide information about genetic relationships and individual ancestry. With GBS data, patterns of allele frequency differentiation and nucleotide sequence divergence can be compared between populations across a broad sample of the genome, and individuals with hybrid ancestry can be identified. More ancient processes of isolation of populations, gene flow, and adaptive introgression can be inferred by examining relationships between relative differentiation (*F*_ST_), absolute between-population genetic distances (π_B_, also known as *D*_xy_), and within-population genetic diversity (π_W_) (Cruickshank & Hahn 2014; Irwin et al. 2016, 2018).

The Pacific Wren (*Troglodytes pacificus*) and Winter Wren (*Troglodytes hiemalis*) are two closely-related songbirds that provide an interesting case of a narrow contact zone between two similar-looking species that started diverging in the distant past. These two species were previously considered conspecific along with the Eurasian Wren (*Troglodytes troglodytes*), but in 2010 they were designated as separate species (Chesser et al., 2010) based on maintenance of substantial genetic and song differentiation in an overlap area in northeastern British Columbia (BC), Canada (Toews & Irwin, 2008). The existence of cryptic species in this system had been first suggested by song differentiation between the populations (Kroodsma & Momose, 1991). While extremely variable, songs of Pacific Wrens in western North America were markedly different than those of Winter Wrens in eastern North America, the latter of which was more similar to songs across Eurasia. This pattern was later reflected in mitochondrial DNA sequences, which showed three major wren clades, with Winter Wrens being more similar to Eurasian Wrens than they were to Pacific Wrens (Drovetski et al., 2004). Toews and Irwin (2008) assessed nuclear DNA differentiation by using 90 Amplified Fragment Length Polymorphism (AFLP) markers and these showed strong differentiation between the Mikkelsen & Irwin, 08 Feb 2021 – preprint copy - BioRxiv Pacific and Winter Wrens, both in allopatry and within the narrow contact zone in northeastern BC. One likely first-generation hybrid was identified by this study (Toews & Irwin, 2008), but the limitations of AFLP data made it difficult to confirm this identity with confidence.

While most temperate songbird species pairs have been estimated to have diverged within the last one million years (Weir & Schluter, 2007), the mitochondrial divergence of Pacific and Winter Wren appears remarkably old, estimated at 4.3 million years ago (Toews & Irwin, 2008) using a standard molecular clock of 2% sequence divergence per million years (Weir & Schluter, 2008). These wrens provide an intriguing opportunity to glimpse the genomic patterns of differentiation in an older species pair with greater reproductive isolation than many recently-studied bird taxa (e.g., Morales et al., 2016; Baldassarre, White, Karubian, & Webster, 2014; Toews et al., 2016; Brelsford & Irwin 2009).

In this study we use genotyping-by-sequencing to assess evidence for differentiation, hybridization, and introgression across the genomes of Pacific and Winter Wrens, from regions of both allopatry and sympatry in North America (Figure 1A). Using variation at hundreds of thousands of SNPs distributed through the genome, we ask: 1) How differentiated are Pacific and Winter Wrens, in terms of both relative differentiation and genetic distance?; 2) Is there hybridization between the species, and what classes of hybrids are present (e.g., F1, F2, backcross)?; and 3) Is there evidence for past introgression between the species? We use answers to these questions to infer the relative importance of assortative mating and low hybrid fitness in speciation of these wrens. Our findings provide substantial insight into why there is not more overlap in breeding range between the two species. These conclusions are likely relevant to the dynamics of many contact zones between closely related species.

**Figure 1.**
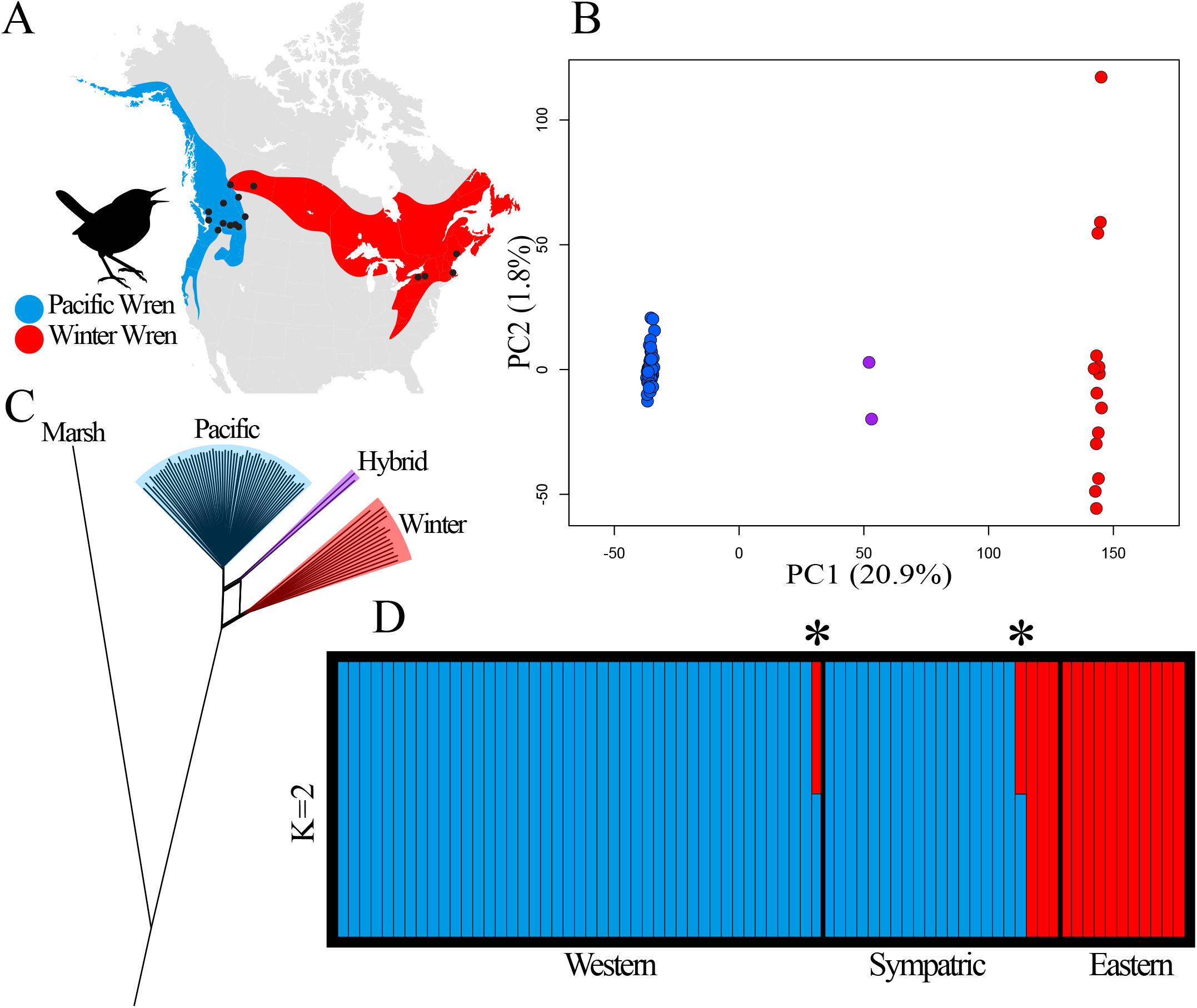
Genetic differentiation of Pacific and Winter Wrens. **A:** range of the Pacific (blue) and Winter (red) Wrens in North America. Sampling localities are indicated by black dots. **B**: Principal Components Analysis. Along the first major axis of variation (PC1), two distinct clusters correspond to Pacific (blue) and Winter Wrens (red), with two intermediate individuals inferred to be hybrids (purple). **C**: Phylogenetic network showing two clades of Pacific (blue) and Winter Wrens (red). The two hybrids (purple) are placed at an intermediate position with ancestry from both parental clades. A Marsh Wren (*Cistothorus palustris*) forms the outgroup. **D**: STRUCTURE analysis with *K* = 2 populations assigns proportions of each sample to two populations, which correspond to Pacific (blue) or Winter Wren (red). All non-hybrid samples show 100% assignment to one population. Two hybrids (asterisks) show intermediate population assignment between Pacific and Winter Wren, with 52% and 48% assignment respectively, which together with high heterozygosity is indicative of them being F1 hybrids.

While it was not the primary purpose of this study, we additionally demonstrate how comparisons of GBS data between females and males can be used to identify interchromosomal translocations between autosomes and sex chromosomes, and we document one of the few known examples in birds.

## Methods

### Sample Collection

We used DNA samples from a total of 76 Pacific and Winter Wrens in this study, including 43 wrens from the allopatric range of the Pacific Wren, 22 from sympatry in Tumbler Ridge BC, and 11 from the allopatric range of the Winter Wren. Sampling locations are summarized in Table S1, and information from each individual can be found in Table S2. Most samples analyzed in this study were collected previously for AFLP analysis by Toews & Irwin (2008). Wrens were attracted by song playback and temporarily captured using mist-nets, and blood samples were taken from the brachial vein and stored in Queen’s Lysis Buffer (Seutin, White, & Boag, 1991). A Marsh Wren (*Cistothorus palustris*) from Vancouver, BC was included for use as an outgroup. *Cistothorus* is thought to belong to the sister group of the Winter Wren complex (Barker, 2017; Gomez, Barber, & Peterson, 2005; Mann, Barker, Graves, Dingess-Mann, & Slater, 2006), with an estimated divergence time of 5 million years ago (Barker, 2017).

### GBS Library Preparation

DNA samples were extracted from whole blood or tissues using a standard phenol-chloroform extraction procedure. Original DNA extracts previously prepared for AFLP analysis (Toews & Irwin, 2008) were available for most samples, and these were supplemented by re-extracted DNA when required to achieve 100 ng DNA per sample. We measured DNA concentrations with a Qubit fluorometer (Invitrogen), and diluted or re-concentrated samples to 20 ng/μL in TE buffer.

Samples were genotyped using genotyping-by-sequencing (GBS), described by Elshire et al. (2011) with modifications by Alcaide, Scordato, Price, and Irwin (2014). This method is based on sequencing a library of size-selected amplified DNA fragments that are from sequences adjacent to restriction enzyme recognition sites throughout the genome. Fragments from each individual are labelled with a unique barcode, allowing them to be pooled, sequenced, and then sorted by individual. This method can produce a dataset of many thousands of SNPs throughout the genome.

We digested 100 ng of DNA of each sample using PstI restriction endonuclease (New England Biolabs), and then ligated fragment ends to common and barcode adapters. A no-barcode blank and a no-DNA blank were run alongside the samples as negative controls. Samples were purified using AMPure XP Beads (Beckman-Coulter) at a ratio of 23:15 in order to eliminate fragments smaller than 100 bp. We amplified the fragments via PCR with Phusion High-fidelity Taq polymerase (New England Biolabs), with the following protocol: 98°C (30 sec) followed by 18 cycles of 98°C (10 sec), 65°C (30 sec), and 72°C (30 sec), with a final extension step at 72°C (5 min). See Figure S1 for primer sequences. PCR was performed separately on each sample prior to pooling in order to achieve more uniform coverage of each sample in the pooled library. Concentrations of PCR products were fluorometrically assayed, and the library was pooled with 100 ng of each sample and control. The DNA was then concentrated in a vacuum centrifuge. We ran the library in three lanes of a 2% agarose gel and extracted the 300-400 bp fragments using a QIAquick Gel Extraction Kit (Qiagen) following the manufacturer’s instructions. The final library was diluted to a concentration of 2 ng/μL in 50 μL. The size range of the fragments was examined using a Bioanalyzer, and the library was sequenced using one lane of Illumina HiSeq 2500 at the Genome Quebec Innovation Centre. This generated paired-end 125 bp reads.

### Bioinformatics Pipeline

We demultiplexed and removed barcode and adapter sequences according to the methods used by Irwin, Alcaide, Delmore, Irwin, and Owens (2016), allowing no barcode sequence mismatches. Reads were trimmed using TRIMMOMATIC v0.32 (Bolger, Lohse, & Usadel, 2014) with the following settings: Trailing:3, Slidingwindow:4:10, and Minlen:30. This trims the 3’ end of reads where average base quality drops below 10 within a four-base sliding window, removes trailing bases below a quality of 3, and retains reads at least 30 bases in length. Reads were mapped to the *Ficedula albicollis* reference genome version 1.5 (Ellegren et al., 2012) using BWA-MEM v0.7.17 (Li & Durbin, 2009), and the resulting SAM files were converted to BAM format using Picard v1.97 (http://broadinstitute.github.io/picard/). The single-end and paired-end BAM files were merged using Samtools (Li et al., 2009). Single nucleotide polymorphisms (SNPs) were identified using GATK v4.0.1.0 (McKenna et al., 2010) HaplotypeCaller, with max alternate alleles set to 2. We used VCFtools (Danecek et al., 2011) to remove indels, SNPs with more than two alleles, SNPs with more than 30% missing genotypes (max-missing=0.7), sites with mapping quality (MQ) below 20.0, and sites with heterozygosity above 60%. This produced a GVCF file for each chromosome containing the genotypes of each individual.

### Principal Components Analysis

We used Principal Components Analysis (PCA) to visualize patterns of genetic clustering in the genotype dataset in *R* (v1.1.383) (R Core Team, 2018) with custom R scripts by Irwin et al. (2016, 2018) using the package PCAMETHODS to impute missing data with svdImpute (Stacklies, Redestig, Scholz, Walther, & Selbig, 2007). We excluded loci where the minor allele was observed only once (i.e., minor allele count < 2) and then conducted PCA for each chromosome individually. A closely-related pair of *T. pacificus* individuals was detected and one individual of the pair was excluded (Table S2) to avoid the PCA being heavily influenced by that close relationship. PCA was then repeated for Pacific Wrens and Winter Wrens separately to investigate population structure within each species, and to investigate if the samples in the range of the subspecies *Troglodytes pacificus salebrosus* in southeastern BC and central Washington could be genetically distinguished from nominate *T. p. pacificus* samples.

### STRUCTURE Analysis

To objectively assign individuals to populations and to further investigate levels of introgression, 20,000 of 127,778 phylogenetically informative SNPs (i.e., SNPs in which the minor allele was observed in more than one individual) were selected to analyze in STRUCTURE (v2.3.4) (Stephens, Pritchard, & Donnelly, 2000). SNPs were randomly selected within each chromosome, with the representation of each chromosome proportional to the number of mapped SNPs it contained. STRUCTURE separates samples into populations by minimizing the linkage and Hardy-Weinberg disequilibrium between alleles within populations, and assigns a proportion of each individual’s genome to each population. Analyses were run with the Admixture Model and assumption of correlated allele frequencies, with 100,000 Markov Chain Monte Carlo steps after a burn-in period of 100,000 steps. Simulations were run with ten replicates each from *K*=1 to *K*=5 populations. The optimal number of populations was identified by Structure Harvester (Earl & vonHoldt, 2012) using the Evanno method (Evanno, Regnaut, & Goudet, 2005), which maximizes the rate of change in the log likelihood of the data (Δ*K*) given a simulated number of populations. Results from each run for the optimal *K* were then combined using the FullSearch algorithm of CLUMPP (v1.1.2) (Jakobsson & Rosenberg, 2007) and plotted using DISTRUCT (v1.1) (Rosenberg, 2004).

### Phylogenetic Network

To examine the phylogenetic relationships and possible patterns of reticulation between lineages, a phylogenetic network was constructed using SplitsTree (v4.14.6) (Huson & Bryant, 2006). This approach was chosen to reflect the possibility of net-like patterns of relatedness within species and between hybridizing lineages, which may not be reflected in a single bifurcating tree, since portions of each individual’s genome will be inherited with recombination from many different ancestors. Distances were calculated using the UncorrectedP method, and a split network was formed using the NeighbourNet method and drawn with the Rooted EqualAngle algorithm. The Marsh Wren sample was used as an outgroup to root the tree.

### Patterns of Differentiation across the Genome

To visualize patterns of differentiation and genetic distance across the genome, we plotted *F*_ST_ (allele frequency differentiation), π_B_ (between-species sequence distance; sometimes referred to as *D*_xy_), and π_W_ (within-species sequence diversity) for sliding windows each containing 10,000 bases of mapped sequences across each chromosome. Note that each 10,000 base window represents a much larger unsequenced distance of the chromosome. Plotting *F*_ST_ allows for the identification of regions with fixed or nearly fixed allele differences between populations; such regions of elevated differentiation can be caused by selective sweeps along with restricted gene flow. Pairwise sequence distances π_W_ and π_B_ are the average proportions of nucleotides that differ between homologous sequences from the same (π_W_) or different (π_B_) populations. This can help to distinguish among several models for the formation of regions of high *F*_ST_ (Cruickshank & Hahn 2014; Irwin et al. 2016, 2018). A pattern of regions of high *F*_ST_ having high π_B_ is consistent with incompatibility loci restricting gene flow in specific regions of the genome. In contrast, high *F*_ST_ regions having low π_B_ is consistent with selective sweeps across hybrid zones, followed by differentiation (Delmore et al. 2015; Irwin et al. 2016, 2018). Calculation of π_W_ and π_B_ incorporated both variant and invariant sites. Plots were created using custom R scripts by Irwin et al. (2016, 2018). *F*_ST_ was calculated using the Weir & Cockerham (1984) method, and the following equations were used:

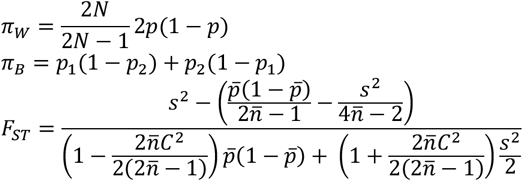

where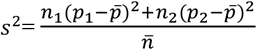 and 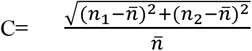 for two populations with allele frequencies *p*_1_ and *p*_2_ and sample sizes *n*_1_ and *n*_2_, with means 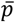 and 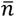.

## Results

### Sequencing results

Sequencing produced 256,467,008 reads totalling 64.12 Gb from 77 individuals (the 76 Pacific or Winter Wrens, and the one Marsh Wren). One sample from Tumbler Ridge generated few reads and was discarded from analysis (Table S2). The filtered sequences altogether mapped to a total of 12.156 Mb of the *Ficedula* genome, which represents 1.2% of the genome assembly (Kawakami et al., 2014). A total of 272,350 SNPs were identified within the Pacific/Winter Wren sample set, for an average of 260 SNPs per megabase. Of these, 144,572 were singletons and 127,778 had minor alleles that were found in more than one individual. An additional 62,819 fixed differences were found between Pacific/Winter Wrens and our Marsh Wren individual (the outgroup), and 681,753 between all of these wrens and the *Ficedula* reference genome. The distribution of SNPs among *Ficedula* chromosomes is summarized in Table S3.

### Principal Components Analysis

Principal components analysis (PCA) revealed two highly distinct genotypic clusters corresponding to Pacific and Winter Wrens widely separated along PC1 (Figure 1B), consistent with previous observations of strong differentiation between the two species in both mitochondrial DNA (Drovetski et al., 2004) and nuclear AFLP markers (Toews & Irwin, 2008). Two individuals, one from Tumbler Ridge BC and one from Gavin Lake BC, were centred between the two clusters on PC1, a pattern expected for F1 hybrids between Pacific and Winter Wrens. The bird from Gavin Lake was previously identified as a possible hybrid by AFLP analysis, although this was considered speculative given the limitations of AFLP data (Toews & Irwin, 2008). Our GBS data showed these two birds repeatedly having intermediate placement on PCAs of each chromosome (Figure S2) and consistent heterozygosity at highly differentiated SNPs along each chromosome, strongly indicating that these are F1 hybrids (later-generation hybrids would show a more variable pattern of inheritance across chromosomes through recombination of parental genotypes). No other intermediate birds appeared on PCAs of any chromosome, indicating that aside from the two F1 hybrids there were no other recent-generation hybrids or backcrosses in our sample. One Winter Wren individual from Tumbler Ridge did fall slightly outside of the Winter Wren cluster on chromosome 24; this individual was heterozygous for a short introgressed region and is discussed further below. For most chromosomes, PC1 separated Pacific and Winter Wrens, while PC2 captured variation within populations; Winter Wrens showed a greater spread of variation along PC2 than Pacific Wrens on most chromosomes, consistent with higher heterozygosity and nucleotide diversity (π_W;_ Table 2), but this was not geographically structured variation.

Principal components analysis within each species revealed no evidence for strong population structure within our sample of either Pacific or Winter Wrens. Pacific Wrens from southeastern British Columbia and central Washington (subspecies *T. pacificus salebrosus*) appeared interspersed with Pacific Wrens of the nominate *Troglodytes pacificus pacificus*, and could not be clearly distinguished by PCA of any chromosome using this dataset (Figure S3A). Similarly, no geographic structure appeared in a PCA of the Winter Wrens, with East Coast birds interspersed with the Albertan and British Columbian samples (Figure S3B).

### STRUCTURE Analysis

STRUCTURE analysis with *K* = 2 populations mirrored the results given by PCA, assigning most individuals to Pacific Wren or Winter Wren clusters with 100% confidence. The only exceptions were the two hybrids, which both showed intermediate proportions of 52% Pacific Wren and 48% Winter Wren (Figure 1D). Structure Harvester identified *K* = 2 populations as optimal for this dataset using the Evanno Method.

### Phylogenetic Network

Pacific and Winter Wrens formed two distinct clades in a phylogenetic network, with shallower divergences within species (Figure 1C). The two hybrids were placed at an intermediate position with ancestry from both the Pacific and Winter Wren clusters. Relative distances between taxa are given in Table 1. Pacific Wrens showed less within-group variation than Winter Wrens, consistent with a narrower spread on the PCA. The distance between the Pacific and Winter Wren clades was 1.38 times wider than the average distance between Pacific Wrens, while the distance between the Marsh Wren and Pacific Wrens was 2.64 times wider than the distance between Pacific and Winter Wrens.

**Table 1.** Relative distances between taxa calculated from a SplitsTree matrix of distances between individuals. Distances were standardized by the average distance between Pacific Wren individuals, and are proportional to the number of SNP differences between individuals. Standard deviations are given in parentheses.

|  | *T. pacificus* | *T. hiemalis* | *C. palustris* |
| --- | --- | --- | --- |
| *T. pacificus* | 1 (0.02) | 1.62 (0.03) | 4.28 (0.01) |
| *T. hiemalis* |  | 1.38 (0.03) | 4.34 (0.02) |

### Limited Introgression Despite Hybridization

To detect historical introgression events that have led to some individuals of one species having genomic segments from the other species, the genotypes of every individual were plotted across each chromosome for all SNPs with *F*_ST_ higher than 0.9 between Pacific and Winter Wrens (Figure S7). A total of 3512 SNPs met this threshold, for an average of 3.4 markers per Mb (note that these are not uniformly distributed, leading to some stretches of lower or higher marker density). At these high-*F*_ST_ loci, Pacific and Winter Wrens are near fixation for different alleles, such that most samples are homozygous for a species-specific allele, while F1 hybrids are heterozygous at most loci. Introgressed regions were identified by eye as stretches of heterozygosity spanning at least five consecutive SNPs in an individual, where nearly all other members of the species are homozygous for a species-specific allele. This method should identify any large introgressed haplotype blocks that span multiple informative SNPs (i.e. >1.5 Mb on average, proportional to SNP density), but would miss very small haplotype blocks that have been broken by many generations of recombination and no longer span multiple informative SNPs. Despite a dataset of many informative SNPs across each chromosome, only one introgressed haplotype block in a single individual was detected (Figure 2, Figure S7), spanning approximately the first 375,000 bp of chromosome 24, corresponding to less than 5 centimorgans on the *Ficedula* genetic map (Kawakami et al., 2014). This individual was a Winter Wren from sympatry near Tumbler Ridge, BC, which contained a Pacific Wren-associated allele at 11 out of 18 high-*F*_ST_ SNPs in this genomic region (other Winter Wrens contained only 0-1 out of 18 Pacific-associated alleles, and the two hybrids contained 9 and 14 out of the 18 Pacific-associated alleles). Given the small size of the remaining *pacificus* haplotype block within this individual, the hybridization event which gave rise to the introgressed region likely ocurred many generations in the past.

**Figure 2.**
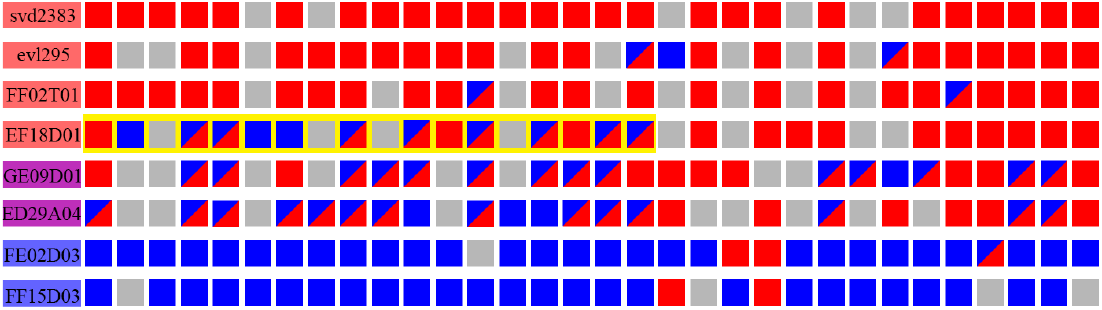
Introgression of a short Pacific Wren haplotype block into a Winter Wren. The first 32 alleles on chromosome 24 with *F*_ST_ greater than 0.9 are plotted for eight representative individuals, showing either the Winter Wren-associated alleles (red blocks), Pacific-associated alleles (blue blocks), heterozygotes (red and blue triangles), or missing data (grey). Winter Wrens are homozygous for the red allele at most loci, while Pacific Wrens are homozygous for the blue allele at most loci and the two F1 hybrids (GE09D01 and ED29A04) are heterozygous at most of the loci. One Winter Wren, EF18D01, contains a Pacific-associated allele at 11 of the first 18 loci (gold highlight), while all other Winter Wrens contain a maximum of 1 Pacific-associated allele at these loci.

### Patterns of Differentiation Across the Genome

To examine patterns of differentiation across the genome, *F*_ST_, π_B_, and π_W_ were plotted along each chromosome in sliding windows that each included a total of 10,000 sequenced basepairs (Figure 3). Differentiation between the species is moderately high across the genome, and most chromosomes show a single broad region of elevated *F*_ST_ above a background of lower *F*_ST._ Mean windowed autosomal *F*_ST_, π_W_ and π_B_ for the three species is given in Table 2. Regions of elevated differentiation tend to have both lower between-population divergence and within-population sequence diversity (Figure 4, Table 2). *F*_ST_ shows a trend toward inverse correlation with between-group pairwise nucleotide distance π_B_ (Figure S4; Pearson’s product-moment correlation, t = −1.94, df = 1159, R = −0.057, p = 0.052). Within-group pairwise nucleotide distance π_W_ is highly inversely correlated with *F*_ST_ (as expected given their mathematical relationship) (Figure S5, Table 2). Regions of increased within-population diversity also tend to have greater sequence divergence between species, as π_W_ and π_B_ are highly correlated (Spearman’s Rank Correlation, S = 3.6 × 10^7^, r_s_ = 0.86, p < 2 × 10^−16^). The windows of highest *F*_ST_ tend to occur at low values of π_B_ and at low values of π_W_ (Figure 4, S4). In summary, we observe moderately high levels of differentiation across the genomes of the Pacific and Winter Wrens, with the regions of highest differentiation occurring in regions of lower sequence diversity (π_W_) and lower absolute sequence divergence (π_B_) (Figure 4, S4, Table 2).

**Table 2.**
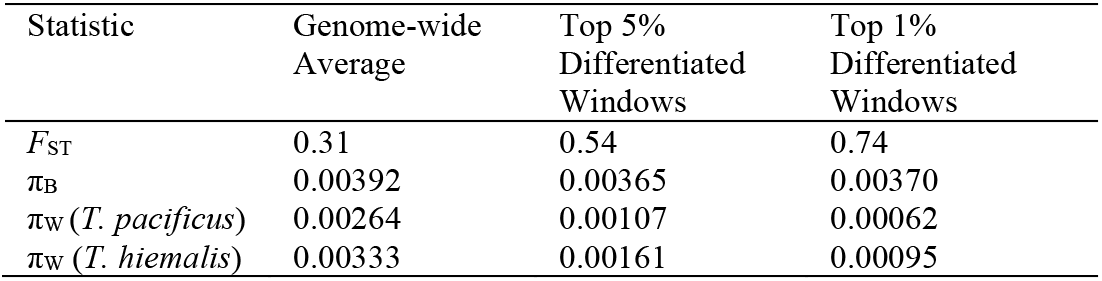
Patterns of differentiation and divergence between Pacific and Winter Wrens. *F*_ST_, π_B_, and π_W_ are averaged across autosomal windows, and are calculated for all windows as well as for the top 5% and 1% of windows with the highest *F*_ST_.

**Figure 3.**
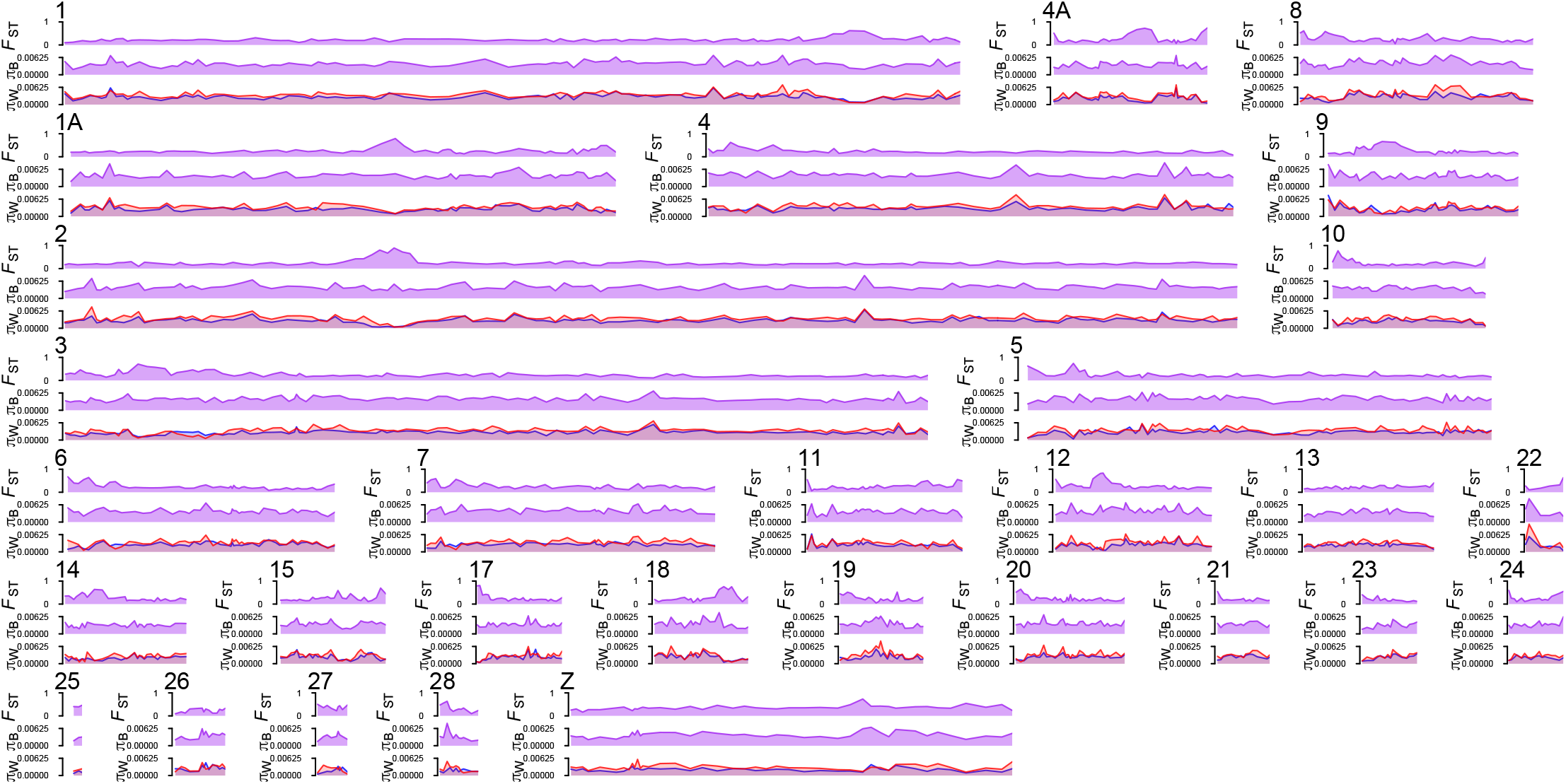
Peaks of divergence and differentiation across the genome. Each chromosome is plotted with windowed *F*_ST_ (allele frequency differentiation, top) and π_B_ (between-species pairwise sequence divergence, middle) between Pacific and Winter Wrens, as well as π_W_ (within-species pairwise sequence diversity, bottom) for Pacific (blue) and Winter Wrens (red).

**Figure 4.**
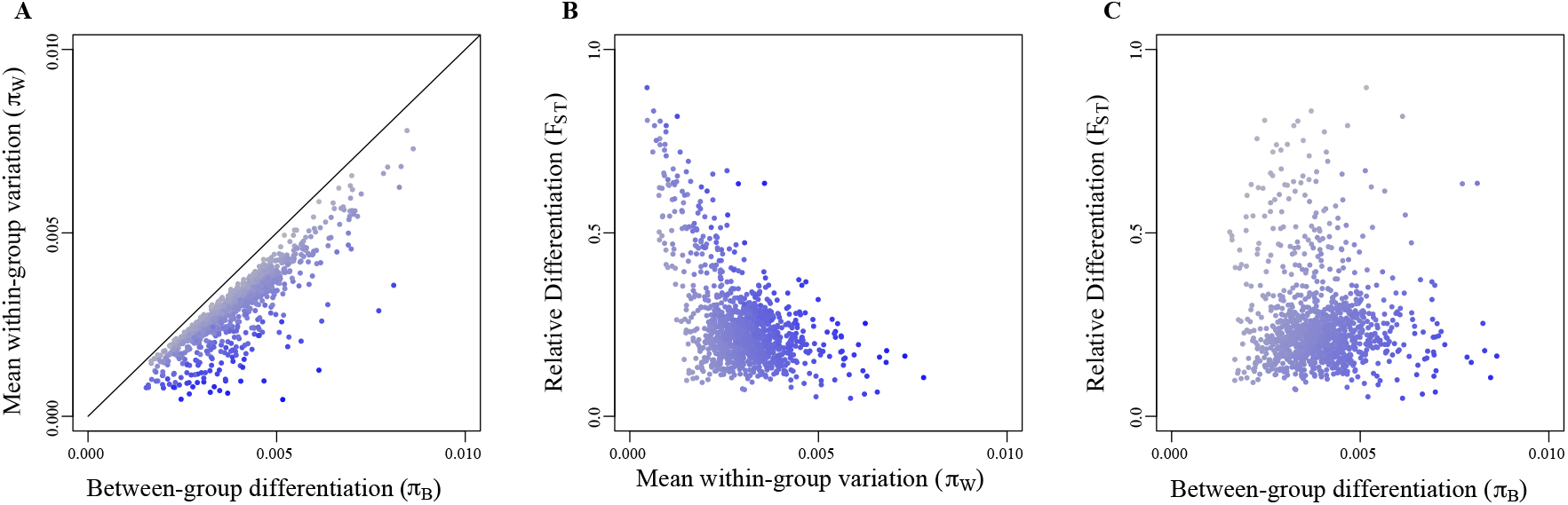
Relationships between π_B_ and mean π_W_ (**A**), mean π_W_ and *F*_ST_ (**B**), and π_B_ and *F*_ST_ (**C**). Statistics were calculated in sliding windows containing 10,000 bp of genotyped positions. Points are colored to depict the unplotted statistic in each panel: *F*_ST_(**A**), π_B_ (**B**), and π_W_ (**C**), with values colored from low (grey) to high (blue).

Differentiation and nucleotide diversity also show a relationship with the type and size of chromosomes (Figure 5). The Z chromosome shows elevated *F*_ST_ relative to the autosomes (0.43 vs 0.31, Figure 5B), and lower π_W_ relative to autosomes of a similar size (Figure 5A). Additionally, both π_W_ and π_B_ are positively correlated with chromosome size, with the steepest relationship among the smaller chromosomes (Figure 5A). These smallest microchromosomes also have the highest *F*_ST_ (Figure 5B, Table S2).

**Figure 5.**
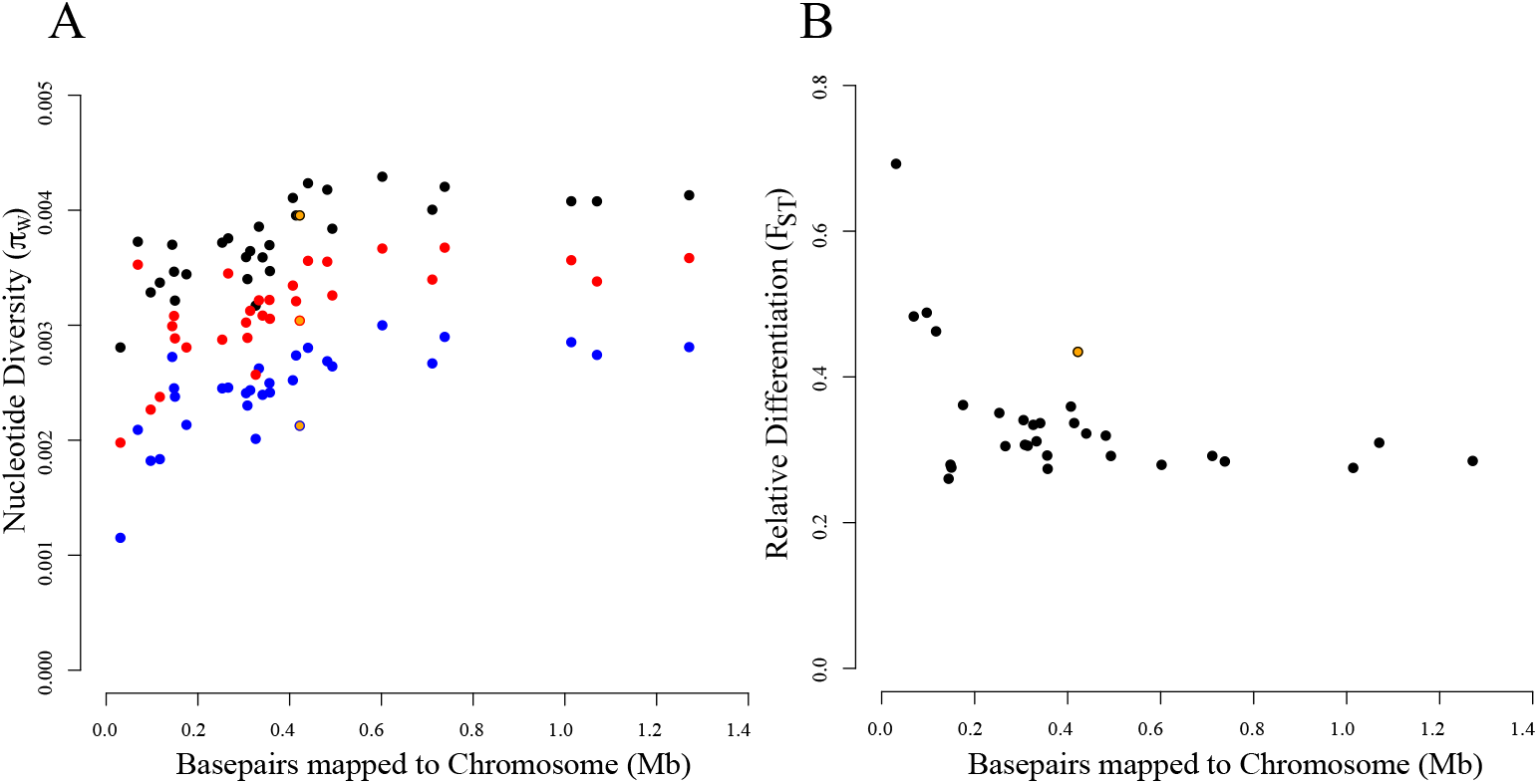
Average *F*_ST_, π_W_, and π_B_ compared to chromosome size. **A**: π_B_ (black) and π_W_ in Pacific (blue) and Winter Wrens (red) show a positive relationship with chromosome size, and the smallest microchromosomes show the lowest levels of nucleotide diversity. Chromosome size is approximated by the number of sequenced basepairs mapped to each *Ficedula* chromosome. The Z chromosome is highlighted in gold. **B**: The four smallest microchromosomes show the highest *F*_ST_, and the Z chromosome (gold) has elevated *F*_ST_ relative to autosomes of a similar size. *F*_ST_ was calculated using the Weir & Cockerham (1984) method for incorporating multiple loci.

### Translocation from an Autosome to a Sex Chromosome

Six females (three Pacific Wrens and three Winter Wrens) were identified in this dataset by observing a substantial reduction in apparent heterozygosity of SNPs mapping to the Z chromosome compared to males (Figure S6) (in birds, females are the heterogametic sex with a Z and W chromosome, while males have two Z chromosomes; females thus only appear heterozygous at the small Z chromosome regions that are homologous with the W chromosome). Surprisingly, these six females were also outliers on a PCA of SNPs mapping to *Ficedula* chromosome 8, being widely separated from all males along PC2 (Figure 6A, Figure S2). The probability that all six of the *Ficedula* chromosome 8 outlier individuals in this dataset would be the six females due to chance is extremely low (*p* = 4.22 × 10^−9^), so it is unlikely that this pattern was not causally related to the sex of the bird. The region of *Ficedula* chromosome 8 that drives this pattern was identified by analyzing successively smaller segments of the chromosome on PCA, which revealed that the boundaries of the region most likely lay between 17.7-17.8 Mb and between 22.2-22.4 Mb of the *Ficedula* chromosome 8 assembly. Since these breakpoints lie near the HS2ST1 and OLFM3 genes, we will refer to this segment of *Ficedula* chromosome 8 as the HS2ST1-OLFM3 region, a roughly 4.5 Mb region containing approximately 74 genes (Table S4).

**Figure 6.**
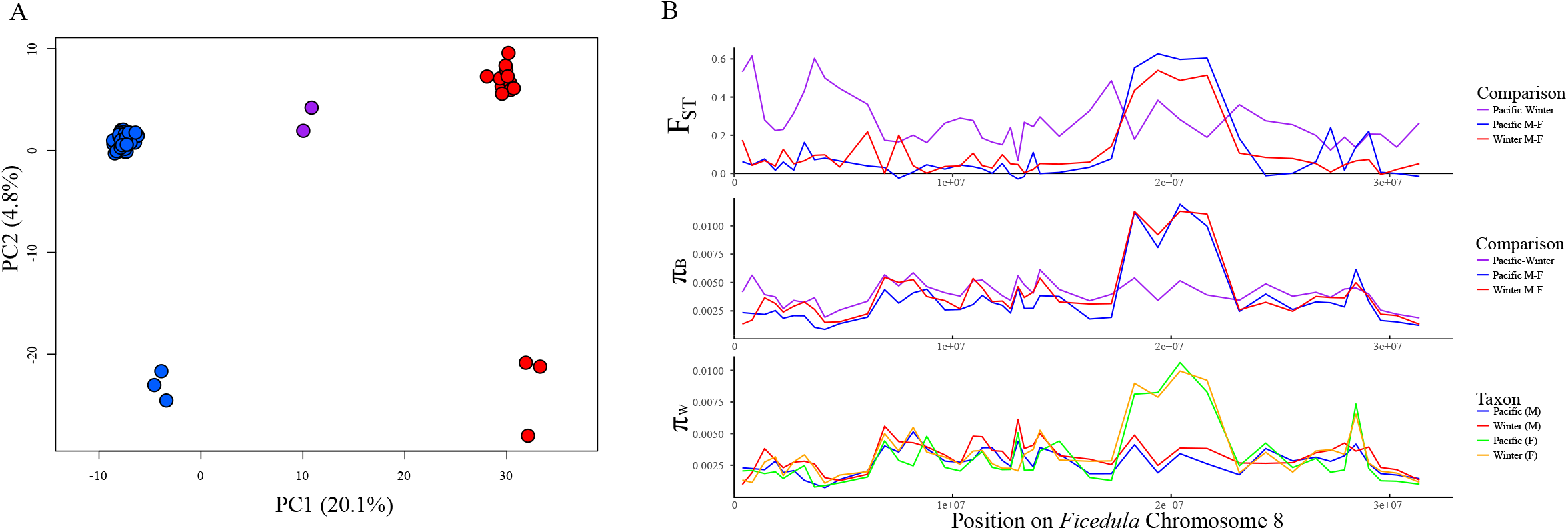
Evidence for a translocation from an autosome to a sex chromosome. **A:** PCA based on sequences mapping to *Ficedula* chromosome 8 separate the six females (three Pacific Wrens, in blue; three Winter Wrens, in red) from the males along PC2, which is not an expected pattern for an autosome. **B:** Scan of *F*_ST_, π_B_, and π_W_ along the sequenced loci mapping to *Ficedula* chromosome 8, with Pacific and Winter Wrens separated into males and females. A large peak appears in the region around 20 Mb in which females have highly elevated π_W_ relative to males (bottom), and where *F*_ST_ (top) and π_B_ (middle) between males and females of the same species is much greater than between males of different species.

The separate clustering of males and females on the PCA of the HS2ST1-OLFM3 region could be explained by a translocation of the region from the autosome to a sex chromosome in the ancestral wren lineage. It is unlikely that the translocation occurred in the opposite direction, from the sex chromosome to chromosome 8 in the *Ficedula* lineage, as gene content in the HS2ST1-OLFM3 and flanking regions of chromosome 8 is conserved with more distant outgroups (Table S4).

The present chromosomal location of the HS2ST1-OLFM3 sequence in the wren genome can be determined by examining patterns of heterozygosity and nucleotide diversity. If the HS2ST1-OLFM3 region were translocated or duplicated to a sex chromosome in the wren lineage, then it would be expected that females and males would show a different level of observed heterozygosity. If the HS2ST1-OLFM3 region were translocated only to the Z chromosome, then males would show a higher apparent heterozygosity than females since males possess two Z chromosomes. Alternatively, if the HS2ST1-OLFM3 region were duplicated to the W chromosome, then females would likely show a higher heterozygosity than males due to female-restricted alleles arising on the W chromosome copy.

To test which predicted pattern is consistent with the data, we compared the apparent heterozygosity of males and females along *Ficedula* chromosome 8. In the region from 17.79-22.30 Mb on *Ficedula* chromosome 8 (the HS2ST1-OLFM3 region), many loci showed much higher heterozygosity in females than males: females are heterozygous at an average of 25% (Pacific) or 28% (Winter) of the SNPs in this region, while males are only heterozygous at 4% (Pacific) or 3% (Winter) of the SNPs (similarly, females showed a much higher value for π_W_ (Table 3, Figure 6B)). The elevated female heterozygosity strongly indicates that a copy of the HS2ST1-OLFM3 region is present in a non-recombining region of the W chromosome, providing allelic diversity that is unique to females. Males still contain sequence reads that map to the HS2ST1-OLFM3 region, indicating that a copy of the sequence either remains on an autosome (chromosome 8) or was also translocated to the Z chromosome. Estimates of sequence diversity provide a way to differentiate between 1) translocation to the W and Z chromosomes with loss of the autosomal copy, 2) translocation to the W and Z chromosomes with the autosomal copy retained, and 3) translocation only to the W and not the Z, with the autosomal copy retained.

**Table 3.** Average π_W_ in males and females in the translocated region homologous to *Ficedula* chromosome 8 from 17.79-22.30 Mb, as well as upstream and downstream sequences of chromosome 8, and the Z chromosome. Females have a higher average π_W_ than males in the translocated region, but comparable π_W_ to males in the chromosome 8 flanking regions and lower π_W_ on the Z chromosome.

|  | Translocated Region | Chr 8 Downstream | Chr 8 Upstream | Chr Z |
| --- | --- | --- | --- | --- |
| Male *pacificus* | 0.0030 | 0.0027 | 0.0027 | 0.0021 |
| Male *hiemalis* | 0.0038 | 0.0033 | 0.0030 | 0.0030 |
| Female *pacificus* | 0.0090 | 0.0024 | 0.0026 | 0.0014 |
| Female *hiemalis* | 0.0092 | 0.0029 | 0.0029 | 0.0022 |

Under the first scenario (translocation to Z and W, loss on autosome), males would likely show a lowered π_W_ in the HS2ST1-OLFM3 region compared to other regions of *Ficedula* chromosome 8 if the translocation event occurred recently, since the translocation event likely involved only a single copy of the sequence lacking any genetic diversity. This pattern is not observed (Figure 6B). However, if this translocation event occurred very far in the past, then subsequent mutations could have restored genetic diversity in the HS2ST1-OLFM3 region. This scenario can be ruled out by comparing the heterozygosity of males and females at loci within the translocated region. If the region were translocated to both the Z and W chromosome and lost on the autosome, then females would show a lower heterozygosity than males at the loci that are variable in males. This is because any new derived alleles that arise on the Z chromosome will have two chances to be found in males with two Z chromosomes, while females will have only one chance to carry the derived Z-linked allele. However, this pattern was not observed: females and males showed similar levels of heterozygosity at the loci that are variable in males (13.8% in females, 13.0% in males).

Under the second scenario (duplication to both the Z and W while retained on an autosome), it would be expected that males would have elevated heterozygosity in the HS2ST1-OLFM3 region relative to the background diversity on *Ficedula* chromosome 8, due to the presence of four rather than two copies of the HS2ST1-OLFM3 region; this is not observed either. Males show a comparable level of heterozygosity within and outside of the translocated region, with males heterozygous at 0.09% (Pacific Wren) and 0.13% (Winter Wren) of all loci within the translocated region, compared to 0.11% and 0.13% of loci in the remaining regions of *Ficedula* chromosome 8. These patterns are instead consistent with the third scenario: that the HS2ST1-OLFM3 region has been duplicated only to the W chromosome while a version of the original autosomal copy has been retained.

## Discussion

In both allopatry and sympatry, Pacific Wrens and Winter Wrens form highly distinct genetic clusters, with an overall genomic *F*_ST_ of 0.31. Together with clear phenotypic differences in song and migratory behavior as well as subtle difference in plumage coloration (Toews & Irwin 2008), the evidence in support of them being separate species is strong. The estimated mitochondrial DNA divergence time indicates that these two species started diverging long ago (roughly 4.3 million years ago) compared to many other sister species of birds (Weir & Schluter, 2004).

Surprisingly given this deep age of divergence between Pacific and Winter Wrens, two first-generation hybrids (i.e., F1 hybrids) were observed within a dataset of 75 wrens, only 22 of which were from the contact zone between the two species. These two observed F1 hybrids were both adult birds who had successfully undergone one cycle of seasonal migration between breeding and wintering grounds, indicating that they are healthy enough to survive these phases of the life cycle. They were found far apart from one another, meaning that they almost certainly are the products of different parental pairs. Hence these F1 hybrids show that premating isolation is far from complete and that F1 hybrids can be healthy.

Despite production and viability of F1 hybrids, we saw no evidence for recent backcrossing or other reproduction of hybrids, suggesting that F1 hybrids suffer greatly reduced fitness relative to parental birds. The most plausible explanation for our results is that F1 hybrids currently have low (virtually zero) reproductive success. Without strong selection against hybrids, later-generation hybrids should outnumber first-generation F1 hybrids given that F1 hybrids are only produced by an initial hybridization event and any of their descendants would be detectable as hybrids for many generations. Determining the reason for low fitness of F1 hybrids will have to await further study, but possibilities include problems with meiosis, behaviors of hybrids that interfere with processes such as seasonal migration (Helbig, 1991; Delmore, Toews, Germain, Owens, & Irwin, 2016) or mate attraction (Bridle, Saldamando, Koning, & Butlin, 2006), and physiological problems that interfere with reproduction or cause the offspring of F1 hybrids to be inviable (Hill, 2019).

Despite the lack of evidence for recent gene flow between Pacific and Winter Wrens, we were able to detect some evidence for gene flow in the more distance past. By examining thousands of markers across the genome, it is possible to detect hybridization events from past generations by observing blocks of markers in the genome containing alleles associated with the other species (Alcaide et al., 2014; Lopes et al., 2016; Ravinet et al., 2017; Sedghifar, Brandvain, & Ralph, 2016; VonHoldt et al., 2011). Only one such block was detected, in a sympatric Winter Wren that is heterozygous for a single 375 kb Pacific Wren haplotype fragment on chromosome 24. The small block size, corresponding to less than 5 cM on the *Ficedula* genetic map (Kawakami et al., 2014), and absence of similar detectable fragments on other chromosomes strongly suggest that this introgression event occurred many generations in the past.

The finding of only a single block of clear introgression between the two species indicates only a very small amount of introgression in the relatively recent past (e.g., on the order of the past hundred to several thousand generations). More deeply in time, examining the relationship between *F*_ST_ and π_B_ can help distinguish between different histories of population isolation and/or gene flow (Cruickshank & Hahn 2014; Irwin et al. 2016, 2018). The lack of a positive relationship between *F*_ST_ and π_B_ leads us to reject a simple model of differentiation with gene flow, in which reproductive incompatibilities protect segments of the genome from gene flow. The slight negative relationship (although not quite significant) suggests a possible role for occasional selective sweeps between the two species, followed by selective differentiation of these regions within each population. Such a history is compatible with the many cycles of glacial and interglacial periods in North America over the past 2 million years, in which incipient Pacific and Winter wren populations could have undergone periods of isolation followed by contact and hybridization. However, the lack of a statistically significant relationship between *F*_ST_ and π_B_ does not allow a clear distinguishing of that model from simple differentiation in allopatry, in which a flat relationship between *F*_ST_ and π_B_ is expected.

Unlike many brightly-plumaged songbirds, the cryptic plumage and subtle morphological differences that distinguish the Pacific and Winter Wrens appear unlikely to present a significant premating barrier between the species. Instead, song, a learned trait, may be critical for species recognition and mate selection as it is in many other bird species (Kroodsma, Bereson, Byers, & Minear, 1989; Mason et al., 2017), and consistent song pattern differences have been identified between the two wren species (Kroodsma, 1980; Toews & Irwin, 2008), although these differences are apparently insufficient to entirely prevent hybridization. Given that a hybrid was found out of only four Winter Wrens (as originally identified in the field) included in our GBS dataset from the sympatric zone, assortative mating may not be strong. Over time, selection against the production of lower-fitness hybrids might drive the evolution of premating barriers (i.e., reinforcement; Dobzhansky 1940; Liou and Price 1994); however, the low density of wrens in the Tumbler Ridge overlap zone may prevent the fixation of assortative mating alleles as these could be swamped by gene flow from regions of allopatry where birds are not subjected to selection against hybridization. With incomplete premating barriers, postmating barriers may be critical in maintaining two distinct species in this system.

Our results provide insight into one curious aspect of the ranges of Pacific and Winter Wrens: both species have large breeding ranges, but only a narrow area of overlap between them. Within and to either side of the area of overlap, both species are found in similar habitat (Toews & Irwin, 2008). Given the results presented here, we propose that the narrow overlap is due primarily to reproductive interference and hybridization between the two species. If two species have only moderate premating isolation but F1 hybrids have zero or very low fitness, then interbreeding in the area of range overlap will result in a large set of offspring that do not themselves successfully reproduce. This can result in population decline, causing a population sink. Movement of the parental forms into the zone can keep the contact zone from disappearing entirely, but it remains an area of low population density. This is an extreme form of the “tension zone” model of hybrid zones, in which low hybrid fitness and dispersal of the parental forms into the zone results in a narrow hybrid zone that is stable for long periods of time (Barton & Hewitt, 1989). The fact that such a pattern (narrow overlap, moderate assortative mating, and very low fitness of F1 hybrids) is seen in such an old species pair suggests that such dynamics explain many narrow hybrid zones.

Compared to strong genetic differences between Pacific and Winter Wrens, we observed little variation within each species. Pacific Wrens from the ranges of subspecies *T. p. pacificus* and *T. p. salebrosus* are not separated by our PCA of this dataset, and only a single SNP showed *F*_ST_ above 0.7 between those groups. This suggests that there is little genomic differentiation between the two subspecies, although it is possible that sampling of wrens from farther southeast in the range of *T. p. salebrosus* would reveal more distinct population structure. Phenotypically, *T. p. pacificus* are slightly darker and more rufescent than *T. p. salebrosus* (Burleigh, 1959), a common trend in birds of the humid Pacific Northwest region. These observed differences may be driven by selection on a few genes above an otherwise homogenous genomic background. If a genetically distinct *T. p. salebrosus* population exists, it may be restricted farther southeast than currently described.

The Z chromosome has a higher average between-species differentiation (*F*_ST_) than autosomes in this dataset, as has been observed in other systems (e.g., Delmore et al., 2015; Oyler-McCance, Cornman, Jones, & Fike, 2015; Corl & Ellegren, 2012; reviewed by Irwin, 2018). This pattern appears to be driven by lower π_W_, rather than higher π_B_, on the Z chromosome relative to autosomes of a similar size (Figure 5A). This lower within-population diversity may be partially attributed to the lower effective population size of the Z compared to autosomes (Corl & Ellegren, 2012; Oyler-McCance et al., 2015), and to the possibility of broader and more frequent selective sweeps due to reduced recombination and exposure of recessive mutations in the heterogametic sex (Borge, Webster, Andersson, & Saetre, 2005).

Interestingly, within-species nucleotide diversity (π_W_) is positively correlated with chromosome size, particularly among the smallest microchromosomes (Figure 5), resulting in the smaller chromosomes having higher average between-species differentiation (*F*_ST_) despite a lower between-species sequence divergence (π_B_). The patterns of π_B_ in the wrens contrast with findings of higher sequence divergence between chicken and turkey microchromosomes than between macrochromosomes (Axelsson, Webster, Smith, Burt, & Ellegren, 2005). The microchromosomes of birds harbour a higher gene density than macrochromosomes (ICGSC, 2004), and it may be that this high gene density has led to stronger and denser background selection or more frequent selective sweeps on the wren microchromosomes, reducing their diversity and leading to faster differentiation. These patterns are consistent with selection on individual loci playing a large role in determining genetic diversity along chromosomes, and they do not match the prediction that microchromosomes should have higher diversity than macrochromosomes based on a positive relationship between recombination rate and sequence diversity (Mugal, Nabholz, & Ellegren, 2013) and a negative relationship between recombination rate and chromosome size (Megens et al., 2009).

Translocations from autosomes to sex chromosomes have been theorized to facilitate sex-specific expression of genes, which may relieve conflict from sexually antagonistic alleles that are beneficial in one sex while detrimental in the other (Albert & Otto, 2005; Irwin, 2018; Rice, 1984). So far, a few examples have been discovered of translocations to the avian sex chromosomes, including in *Myzomela* honeyeaters (Sardell, 2016), *Alauda* larks (Brooke et al., 2010), the Sylvioidea songbird superfamily (Pala et al., 2012), and the Eastern Yellow Robin (Gan et al., 2019). In the present study, the 4.5 Mb region duplicated to the wren W chromosome contains 74 annotated protein-coding genes in the *Ficedula* chromosome 8 assembly. This represents a substantial addition to the 6.9 Mb non-recombining region of the W chromosome, which contains only 46 known genes in *Ficedula* (Smeds et al., 2015). If one of these genes is involved in expressing sexually antagonistic traits, then selection may have favoured a rare duplication of this fragment to the W chromosome despite the apparently strong selection against most inter-chromosomal translocations. It remains to be determined whether these duplicated genes remain functional, and whether divergence in function has occurred between the W and autosomal gene copies. These wrens provide a rare opportunity to investigate the relative rates and characteristics of evolution of homologous autosomal and W-chromosomes copies of a large genetic region.

In conclusion, Pacific and Winter Wrens have been differentiating for millions of years and appear to have almost complete lack of current gene flow. They still hybridize at a sizeable rate, but the F1 hybrids apparently have zero or very low fitness. The combination of little to moderate assortative mating with very low fitness of hybrids is expected to result in populations being unable to coexist over a broad region (Barton & Hewitt, 1989; Goldberg & Lande, 2007; Irwin, 2020). The Pacific and Winter Wrens provide a particularly difficult-to-detect combination of reproductive isolating dynamics: their similar appearances make hybrids difficult to detect, and the fact that F1 hybrids are viable would make low hybrid fitness difficult to detect directly (both of these were inferred here using genomic analysis). We suspect that similar dynamics are common in many other contact zones between closely-related neighbouring species. If so, the narrowness of hybrid zones is explained well by reproductive interference and these tension zone dynamics. While differentiation in ecological niches and/or niche exclusion are interesting factors, in many cases they may not need to be invoked to explain narrow contact zones. Difficult-to-detect production of low-fitness hybrids could provide a large part of the explanation for the general observation that it tends to take many millions of years of differentiation before bird species become broadly sympatric (Price, 2010).

## Acknowledgments

We are grateful to Silu Wang and Armando Geraldes for lab training and support. We also thank Loren Reiseberg, Sally Otto, and Kristin Nurkowski for laboratory support and assistance. We thank David Toews for collecting many of the samples from the Toews and Irwin (2008) study. The Genome Quebec Innovation Centre provided sequencing services. Funding was provided by the Natural Sciences and Engineering Research Council of Canada (grants 311931-2012, RGPIN-2017-03919 and RGPAS-2017-507830 to DEI).

## Author Contributions and Notes

E.K.M. and D.E.I. conceived of the project. E.K.M. performed lab work. D.E.I. and E.K.M. ran the bioinformatics pipeline. E.K.M. analyzed the data and D.E.I. provided guidance. E.K.M and D.E.I. wrote the paper.

## Data Accessibility

Prior to publication, DNA sequencing reads will be deposited to the NCBI Sequence Read Archive. Genotype data and R code associated with the PCA, *F*_ST_, π_W_, and π_B_ analyses will be deposited in Dryad and available in a Github repository (https://github.com/elsemikk/Wren_popgen).

