## Supplemental Information for "Ongoing production of low-fitness hybrids limits range overlap between divergent cryptic species"

**Table of Contents:**

| **Table S1** | Page 2 |
| --- | --- |
| **Table S2** | Page 3 |
| **Figure S1** | Page 5 |
| **Table S3** | Page 6 |
| **Figure S2** | Page 7 |
| **Figure S3** | Page 9 |
| **Figure S4** | Page 10 |
| **Figure S5** | Page 11 |
| **Figure S6** | Page 12 |
| **Table S4** | Page 13 |
| **Figure S7** | Page 17 |

**Table S1.** Sampling locations of individuals analyzed in this study. Allopatric *pacificus* populations are highlighted in blue, sympatric populations in purple, and allopatric *hiemalis* populations in red.

| **Location** | ***pacificus* Samples** | **Hybrid Samples** | ***hiemalis* Samples** | **Latitude** | **Longitude** |
| --- | --- | --- | --- | --- | --- |
| Okanogan, WA | 1 | 0 | 0 | 48.4 | -119.6 |
| Vancouver, BC | 19 | 0 | 0 | 49.3 | -123.2 |
| Whistler, BC | 2 | 0 | 0 | 50.1 | -123.0 |
| Penticton, BC | 2 | 0 | 0 | 49.3 | -119.8 |
| Christina Lake, BC | 2 | 0 | 0 | 49.1 | -118.2 |
| Nelson, BC | 2 | 0 | 0 | 49.6 | -117.3 |
| Cranbrook, BC | 3 | 0 | 0 | 49.4 | -115.9 |
| Hinton, AB | 4 | 0 | 0 | 53.7 | -117.3 |
| Kananaskis, AB | 1 | 0 | 0 | 51.1 | -114.9 |
| Gavin Lake, BC | 6 | 1 | 0 | 52.5 | -121.7 |
| **Total - Allopatric *pacificus*** | **42** | **1** | **0** |  |  |
| Tumbler Ridge, BC | 17 | 1 | 3 | 55.0 | -121.1 |
| Lesser Slave Lake, AB | 0 | 0 | 7 | 55.4 | -115.0 |
| Penobscot, ME | 0 | 0 | 1 | 44.5 | -68.7 |
| Plymouth, MA | 0 | 0 | 1 | 42.0 | -70.7 |
| Dryden, NY | 0 | 0 | 1 | 42.5 | -76.3 |
| Webster, NY | 0 | 0 | 1 | 43.2 | -77.4 |
| **Total - Allopatric *hiemalis*** | **0** | **0** | **11** |  |  |
| **Total** | **59** | **2** | **14** |  |  |

**Table S2**. Metadata for samples used in this project, including PCA and STRUCTURE results.

| Record Number | Species | Sex | Location | PCA_PC1 | PCA_PC2 | PCA_PC3 | STRUCTURE pop1 | STRUCTURE pop2 | Day | Month | Year | Degrees N | Degrees W |
| --- | --- | --- | --- | --- | --- | --- | --- | --- | --- | --- | --- | --- | --- |
| ED29A07 | pacificus | M | Gavin Lake, BC | -35.35764 | 3.21205187 | -15.228351 | 1 | 0 | 29 | 4 | 2005 | 52.50405 | 121.62341 |
| ED26A01 | pacificus | M | Gavin Lake, BC | -35.83976 | 0.24940914 | 27.1750732 | 1 | 0 | 26 | 4 | 2005 | NA | NA |
| ED27A05 | pacificus | M | Gavin Lake, BC | -35.13372 | 4.8377506 | 23.9376409 | 1 | 0 | 27 | 4 | 2005 | 52.48552 | 121.78786 |
| ED28A01 | pacificus | M | Gavin Lake, BC | -33.88786 | 15.6091413 | 1.28677213 | 1 | 0 | 28 | 4 | 2005 | 52.46803 | 121.67396 |
| ED29A02 | pacificus | M | Gavin Lake, BC | -35.36021 | 7.18042982 | -4.3497108 | 1 | 0 | 29 | 4 | 2005 | 52.48146 | 121.69505 |
| ED29A04 | Hybrid | M | Gavin Lake, BC | 53.09487 | -19.844782 | -31.879796 | 0.52 | 0.48 | 29 | 4 | 2005 | 52.4809 | 121.68883 |
| ED29A05 | pacificus | M | Gavin Lake, BC | -35.16653 | 4.61885099 | -4.5791801 | 1 | 0 | 29 | 4 | 2005 | 52.50344 | 121.62505 |
| EE05D01 | hiemalis | M | Lesser Slave Lake, AB | 144.37659 | -1.6432822 | 8.69938063 | 0 | 1 | 5 | 5 | 2005 | 55.48111 | 114.81145 |
| EE05D03 | hiemalis | M | Lesser Slave Lake, AB | 143.26205 | 5.50758095 | 57.1419 | 0 | 1 | 5 | 5 | 2005 | 55.47223 | 114.78301 |
| EE06D01 | hiemalis | M | Lesser Slave Lake, AB | 145.2499 | -15.407428 | 18.440445 | 0 | 1 | 6 | 5 | 2005 | 55.39054 | 114.74566 |
| EE06D02 | hiemalis | M | Lesser Slave Lake, AB | 143.15663 | -29.75394 | -20.778276 | 0 | 1 | 6 | 5 | 2005 | 55.39502 | 114.74916 |
| EE10D01 | failed_GBS | U | Tumbler Ridge, BC | Discarded | Discarded | Discarded | Discarded | Discarded | 10 | 5 | 2005 | 54.85466 | 121.24106 |
| EE25D03 | pacificus | M | Whistler, BC | -36.70272 | -12.659409 | -6.4949385 | 1 | 0 | 25 | 5 | 2005 | 50.06126 | 123.02701 |
| EE29D02 | pacificus | M | Whistler, BC | -35.65012 | -2.2193532 | -9.7067542 | 1 | 0 | 29 | 5 | 2005 | 50.04337 | 123.03195 |
| EF18D01 | hiemalis | M | Tumbler Ridge, BC | 143.36042 | -9.4559996 | -23.688134 | 0 | 1 | 18 | 6 | 2005 | 55.16077 | 120.9472 |
| EF18D03 | pacificus | M | Tumbler Ridge, BC | -35.44146 | -0.4762395 | 14.9625668 | 1 | 0 | 18 | 6 | 2005 | 55.15749 | 120.94101 |
| EF18D04 | hiemalis | M | Tumbler Ridge, BC | 144.23088 | 1.09108973 | 57.0929038 | 0 | 1 | 18 | 6 | 2005 | 55.15555 | 120.94153 |
| EF22D01 | pacificus | M | Tumbler Ridge, BC | -35.54477 | -0.4817343 | 3.47283228 | 1 | 0 | 22 | 6 | 2005 | 55.18352 | 120.9078 |
| EF23D01 | pacificus | M | Tumbler Ridge, BC | -36.43297 | 0.52085327 | -0.0826762 | 1 | 0 | 23 | 6 | 2005 | 54.90497 | 121.22387 |
| EH01D01 | pacificus | M | Pacific Spirit Park, BC | -34.64109 | -2.2766367 | 11.7263717 | 1 | 0 | 1 | 8 | 2005 | 49.25665 | 123.23027 |
| EH03D01 | pacificus | M | Pacific Spirit Park, BC | Excluded | Excluded | Excluded | 1 | 0 | 3 | 8 | 2005 | 49.25301 | 123.22815 |
| EH03D02 | pacificus | M | Pacific Spirit Park, BC | -35.55743 | 0.39711956 | -14.132853 | 1 | 0 | 3 | 8 | 2005 | 49.25682 | 123.23041 |
| EJ29A01 | pacificus | F | Pacific Spirit Park, BC | -35.33918 | 20.6935168 | -22.41963 | 1 | 0 | 29 | 10 | 2005 | 49.25682 | 123.23067 |
| EJ29A02 | pacificus | M | Pacific Spirit Park, BC | -35.35639 | -0.4198556 | 12.4633782 | 1 | 0 | 29 | 10 | 2005 | 49.25682 | 123.23067 |
| EJ29A04 | pacificus | M | Pacific Spirit Park, BC | -36.81872 | -0.802984 | 12.8262734 | 1 | 0 | 29 | 10 | 2005 | 49.25682 | 123.23067 |
| EJ29A05 | pacificus | M | Pacific Spirit Park, BC | -35.96262 | 9.78207434 | 15.5661587 | 1 | 0 | 29 | 10 | 2005 | 49.25645 | 123.23021 |
| EJ29A06 | pacificus | M | Pacific Spirit Park, BC | -35.09761 | 7.10079905 | 20.2295063 | 1 | 0 | 29 | 10 | 2005 | 49.25645 | 123.23021 |
| EK11A01 | pacificus | M | Pacific Spirit Park, BC | -35.34626 | -6.2040677 | 0.05315008 | 1 | 0 | 11 | 11 | 2005 | 49.25529 | 123.22947 |
| EK13A02 | pacificus | F | Pacific Spirit Park, BC | -34.56499 | 20.2331672 | -15.06203 | 1 | 0 | 11 | 11 | 2005 | 49.25401 | 123.22914 |
| EK13A03 | pacificus | F | Pacific Spirit Park, BC | -35.47378 | 11.8861337 | -49.228021 | 1 | 0 | 11 | 11 | 2005 | 49.25401 | 123.22914 |
| FC18D01 | pacificus | M | Pacific Spirit Park, BC | -35.3056 | -3.1756583 | -1.7105506 | 1 | 0 | 18 | 3 | 2006 | 49.272041 | 123.255637 |
| FD17D01 | pacificus | M | Pacific Spirit Park, BC | -35.76917 | 3.43409176 | 18.0305671 | 1 | 0 | 17 | 4 | 2006 | 49.25266 | 123.20561 |
| FD24D01 | pacificus | M | Pacific Spirit Park, BC | -35.95602 | 1.28276896 | 5.27788411 | 1 | 0 | 24 | 4 | 2006 | 49.25632 | 123.23061 |
| FD26D01 | pacificus | M | Pacific Spirit Park, BC | -35.62461 | -5.0054679 | 7.66682829 | 1 | 0 | 26 | 4 | 2006 | 49.25657 | 123.22533 |
| FD26D03 | pacificus | M | Pacific Spirit Park, BC | -35.81045 | -3.650484 | -6.4183526 | 1 | 0 | 26 | 4 | 2006 | 49.25752 | 123.22653 |
| FD28D01 | pacificus | M | Pacific Spirit Park, BC | -36.392 | -1.4270713 | -1.1773148 | 1 | 0 | 28 | 4 | 2006 | 49.27043 | 123.22861 |
| FE02D03 | pacificus | M | Pacific Spirit Park, BC | -35.62868 | 1.73468817 | -0.9508377 | 1 | 0 | 2 | 5 | 2006 | 49.27143 | 123.22593 |
| FE13T02 | pacificus | M | Kananaskis, AB | -36.82614 | -10.030487 | -6.2407218 | 1 | 0 | 13 | 5 | 2006 | 51.05054 | 114.92296 |
| FE19T01 | pacificus | M | Tumbler Ridge, BC | -34.60487 | -1.769967 | -34.929493 | 1 | 0 | 19 | 5 | 2006 | 54.90032 | 121.22266 |
| FE19T02 | pacificus | M | Tumbler Ridge, BC | -34.16362 | 0.84774692 | 7.05540894 | 1 | 0 | 19 | 5 | 2006 | 54.85904 | 121.24375 |
| FE20T01 | pacificus | M | Tumbler Ridge, BC | -35.90848 | 1.28942553 | 9.73473147 | 1 | 0 | 20 | 5 | 2006 | 54.85454 | 121.24117 |
| FE20T02 | pacificus | M | Tumbler Ridge, BC | -35.03914 | -3.5197733 | -0.1226983 | 1 | 0 | 20 | 5 | 2006 | 54.90116 | 121.22386 |
| FE20T03 | pacificus | M | Tumbler Ridge, BC | -35.20132 | -1.2885821 | 4.65567831 | 1 | 0 | 20 | 5 | 2006 | 54.90503 | 121.22398 |
| FE22T01 | pacificus | M | Tumbler Ridge, BC | -36.36269 | 0.64529266 | -4.6554506 | 1 | 0 | 22 | 5 | 2006 | 55.14907 | 120.94897 |
| FE24T01 | pacificus | M | Tumbler Ridge, BC | -35.6146 | -8.8779743 | -7.3931384 | 1 | 0 | 24 | 5 | 2006 | 55.15409 | 120.90332 |
| FE24T02 | pacificus | M | Tumbler Ridge, BC | -35.83875 | -4.9129518 | -7.849324 | 1 | 0 | 24 | 5 | 2006 | 55.15459 | 120.94131 |
| FE25T01 | pacificus | M | Tumbler Ridge, BC | -36.11689 | 0.28217703 | -7.8638737 | 1 | 0 | 25 | 5 | 2006 | 55.10872 | 120.97825 |
| FE27D01 | hiemalis | M | Tumbler Ridge, BC | 142.26996 | 0.31675811 | 4.46183358 | 0 | 1 | 27 | 5 | 2006 | 55.17559 | 120.85533 |
| FF02T01 | hiemalis | M | Lesser Slave Lake, AB | 143.94977 | -25.318076 | 43.5530744 | 0 | 1 | 2 | 6 | 2006 | 55.39579 | 114.25156 |
| FF07T01 | pacificus | M | Hinton, AB | -37.08128 | -3.3486502 | 1.27124914 | 1 | 0 | 7 | 6 | 2006 | 53.70419 | 117.16029 |
| FF08T01 | pacificus | M | Hinton, AB | -35.73332 | 1.46950825 | 0.44299232 | 1 | 0 | 8 | 6 | 2006 | 53.76677 | 117.27464 |
| FF08T02 | pacificus | M | Hinton, AB | -35.99699 | -2.3969143 | -4.3707392 | 1 | 0 | 8 | 6 | 2006 | 53.76562 | 117.29633 |
| FF09T01 | pacificus | M | Hinton, AB | -36.21762 | -0.1522466 | -7.7927779 | 1 | 0 | 9 | 6 | 2006 | 53.70618 | 117.29958 |
| FF11T01 | hiemalis | M | Lesser Slave Lake, AB | 143.18721 | -55.665619 | -34.637086 | 0 | 1 | 11 | 6 | 2006 | 55.51839 | 116.11929 |
| FF11T02 | hiemalis | M | Lesser Slave Lake, AB | 142.7758 | -48.822081 | -22.994853 | 0 | 1 | 11 | 6 | 2006 | 55.51839 | 116.11929 |
| FF13D01 | pacificus | M | Penticton, BC | -34.42139 | 4.59128026 | 6.14697709 | 1 | 0 | 13 | 6 | 2006 | 49.334101 | 119.76691 |
| FF13D02 | pacificus | M | Penticton, BC | -35.42442 | 8.93803483 | 7.76509922 | 1 | 0 | 13 | 6 | 2006 | 49.30882 | 119.78853 |
| FF15D01 | pacificus | M | Christina Lake, BC | -36.69815 | -5.0227528 | 16.8818124 | 1 | 0 | 15 | 6 | 2006 | 49.07296 | 118.16175 |
| FF15D03 | pacificus | M | Christina Lake, BC | -35.3782 | -2.5100563 | 4.52872419 | 1 | 0 | 15 | 6 | 2006 | 49.04493 | 118.25836 |
| FF21T01 | pacificus | M | Cranbrook, BC | -35.46165 | -3.1903707 | -2.8118457 | 1 | 0 | 21 | 6 | 2006 | 49.4331 | 115.91362 |
| FF21T03 | pacificus | M | Cranbrook, BC | -36.18645 | -4.8477895 | -2.4513903 | 1 | 0 | 21 | 6 | 2006 | 49.4277 | 115.92744 |
| FF22T01 | pacificus | M | Cranbrook, BC | -35.89033 | -0.9929037 | 3.81342576 | 1 | 0 | 22 | 6 | 2006 | 49.36302 | 115.55123 |
| FF22T02 | pacificus | M | Nelson, BC | -34.90729 | -0.0396429 | -1.9842591 | 1 | 0 | 22 | 6 | 2006 | 49.5989 | 117.12896 |
| FF23T01 | pacificus | M | Nelson, BC | -35.56324 | 4.12432158 | 15.1565771 | 1 | 0 | 23 | 6 | 2006 | 49.52061 | 117.41111 |
| GD22T01 | C_palustris | M | Pacific Spirit Park, BC | NA | NA | NA | NA | NA | 22 | 4 | 2007 | 49.23706 | 123.2213 |
| GE05D01 | pacificus | M | Pacific Spirit Park, BC | -35.20899 | 4.11165136 | 24.9976945 | 1 | 0 | 5 | 5 | 2007 | NA | NA |
| GE09D01 | Hybrid | M | Tumbler Ridge, BC | 52.0822 | 2.86476671 | -1.9499822 | 0.52 | 0.48 | 9 | 5 | 2007 | 54.861667 | 121.243017 |
| GE11D01 | pacificus | M | Tumbler Ridge, BC | -36.3099 | -2.281343 | 4.74356832 | 1 | 0 | 11 | 5 | 2007 | 54.85568 | 121.2422 |
| GE11D04 | pacificus | M | Tumbler Ridge, BC | -36.43828 | -0.9697357 | 12.1362455 | 1 | 0 | 11 | 5 | 2007 | 54.85443 | 121.24051 |
| GE11D05 | pacificus | M | Tumbler Ridge, BC | -35.58206 | -5.762371 | 9.6737502 | 1 | 0 | 11 | 5 | 2007 | 54.86096 | 121.24318 |
| GE13D01 | pacificus | M | Tumbler Ridge, BC | -35.60595 | -6.6226881 | -19.247895 | 1 | 0 | 13 | 5 | 2007 | 55.18364 | 120.90721 |
| GE14D01 | pacificus | M | Tumbler Ridge, BC | -34.61898 | -6.9087805 | -15.383669 | 1 | 0 | 14 | 5 | 2007 | 55.16934 | 120.96763 |
| evl295 | hiemalis | F | Plymouth, MA | 144.78307 | 59.0389002 | -33.881779 | 0 | 1 | NA | NA | NA | NA | NA |
| sar7443 | hiemalis | M | Penobscot, MN | 144.03151 | -43.68878 | 4.17775412 | 0 | 1 | NA | NA | NA | NA | NA |
| svd2195 | pacificus | M | Okanagan, WA | -36.03893 | -7.293803 | 0.99163149 | 1 | 0 | NA | NA | NA | NA | NA |
| svd2382 | hiemalis | F | Dryden, NY | 143.79005 | 54.6098156 | -55.33097 | 0 | 1 | NA | NA | NA | NA | NA |
| svd2383 | hiemalis | F | Webster, NY | 145.10365 | 117.179208 | 17.4192743 | 0 | 1 | NA | NA | NA | NA | NA |

**GBS-PrimerA:**

5’- AATGATACGGCGACCACCGAGATCTACACTCTTTCCCTACACGACGCTCTTCCGATCT-3’

**GBS-PrimerB:**

5’-CAAGCAGAAGACGGCATACGAGATCGGTCTCGGCATTCCTGCTGAACCGCTCTTCCGATCT-3’

**Figure S1**. Primer sequences used for amplification of GBS library fragments.

**Table S3**. Number of SNPs mapping to each chromosome of *Ficedula albicollis*, as well as mean values for *F*_ST_, π_B_, and π_W_ on each chromosome. Sizes of the chromosomes are reported by Kawakami et al. (2014).

| Chromosome | Size in *Ficedula* (Mb) | Number of SNPs | Total Mapped Sequence (Mb) | Mean *F*_ST_ | Mean π_B_ | Mean π_W_ |
| --- | --- | --- | --- | --- | --- | --- |
| 1 | 119.8 | 24 581 | 1.014 | 0.086 | 0.0041 | 0.0032 |
| 1A | 74.8 | 16 551 | 0.711 | 0.088 | 0.004 | 0.0030 |
| 2 | 157.4 | 30 794 | 1.271 | 0.089 | 0.0041 | 0.0032 |
| 3 | 115.7 | 25 809 | 1.07 | 0.088 | 0.0041 | 0.0031 |
| 4 | 70.3 | 14 516 | 0.602 | 0.088 | 0.0043 | 0.0033 |
| 4A | 21.2 | 6915 | 0.341 | 0.081 | 0.0036 | 0.0027 |
| 5 | 64.6 | 17 622 | 0.738 | 0.083 | 0.0042 | 0.0033 |
| 6 | 37.2 | 11 319 | 0.493 | 0.079 | 0.0038 | 0.0030 |
| 7 | 39.3 | 11 405 | 0.482 | 0.091 | 0.0042 | 0.0031 |
| 8 | 32.0 | 10 246 | 0.44 | 0.087 | 0.0042 | 0.0032 |
| 9 | 26.8 | 8933 | 0.414 | 0.080 | 0.0040 | 0.0030 |
| 10 | 21.3 | 7084 | 0.314 | 0.078 | 0.0036 | 0.0028 |
| 11 | 21.7 | 7554 | 0.333 | 0.080 | 0.0039 | 0.0029 |
| 12 | 21.9 | 9068 | 0.407 | 0.088 | 0.0041 | 0.0029 |
| 13 | 18.6 | 8091 | 0.357 | 0.077 | 0.0035 | 0.0027 |
| 14 | 17.4 | 6079 | 0.326 | 0.076 | 0.0032 | 0.0023 |
| 15 | 14.9 | 6255 | 0.305 | 0.075 | 0.0036 | 0.0027 |
| 17 | 12.4 | 6484 | 0.308 | 0.076 | 0.0034 | 0.0026 |
| 18 | 13.1 | 4774 | 0.253 | 0.082 | 0.0037 | 0.0027 |
| 19 | 11.9 | 5585 | 0.266 | 0.077 | 0.0038 | 0.0030 |
| 20 | 15.6 | 7911 | 0.356 | 0.077 | 0.0037 | 0.0029 |
| 21 | 8.1 | 2970 | 0.148 | 0.075 | 0.0035 | 0.0028 |
| 22 | 5.7 | 944 | 0.069 | 0.107 | 0.0037 | 0.0028 |
| 23 | 7.9 | 2579 | 0.15 | 0.065 | 0.0032 | 0.0026 |
| 24 | 8.0 | 3461 | 0.175 | 0.079 | 0.0034 | 0.0025 |
| 25 | 1.3 | 297 | 0.031 | 0.129 | 0.0028 | 0.0016 |
| 26 | 4.9 | 2815 | 0.144 | 0.081 | 0.0037 | 0.0029 |
| 27 | 4.6 | 1665 | 0.097 | 0.094 | 0.0033 | 0.0020 |
| 28 | 5.0 | 2005 | 0.117 | 0.086 | 0.0034 | 0.0021 |
| Z | 74.6 | 8036 | 0.422 | 0.110 | 0.0041 | 0.0032 |
| **Total:** | **1048** | **272 350** | **12.156** | **0.084** | **0.0038** | **0.0029** |

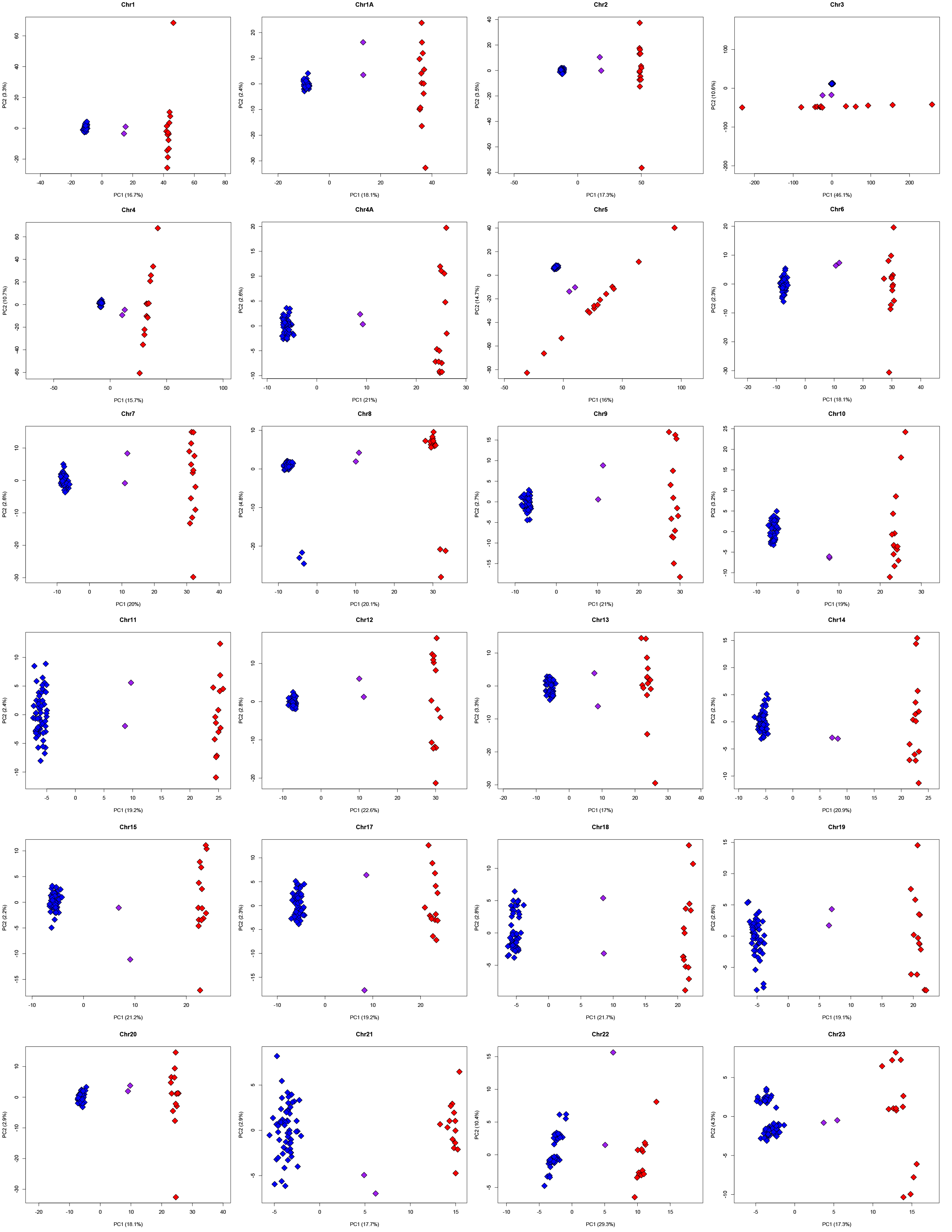

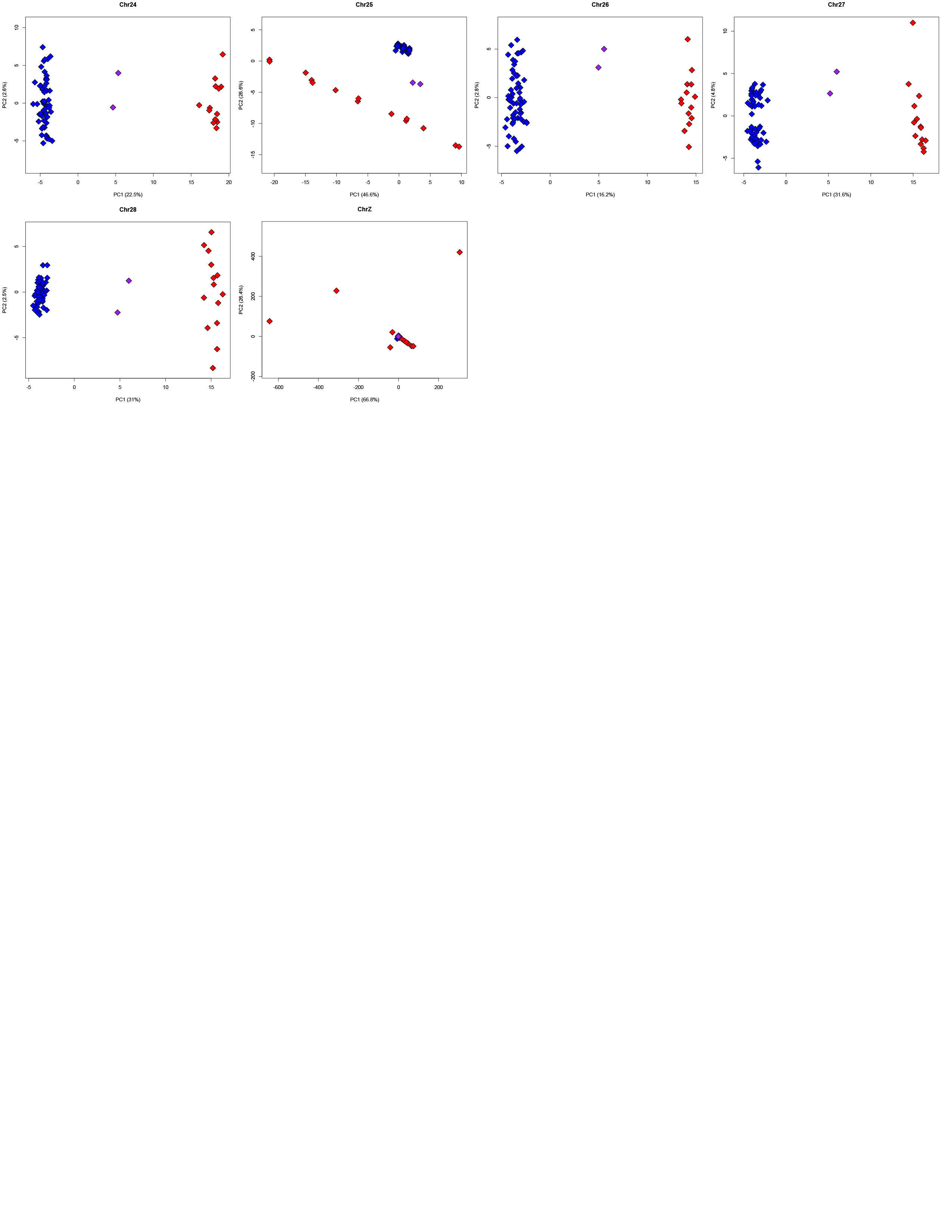

**Figure S2**. PCA analysis of SNPs mapping to twenty-nine autosomes and the Z chromosome of *Ficedula albicollis,* showing Pacific Wrens (blue), Winter Wrens (red), and two first-generation hybrids (purple).

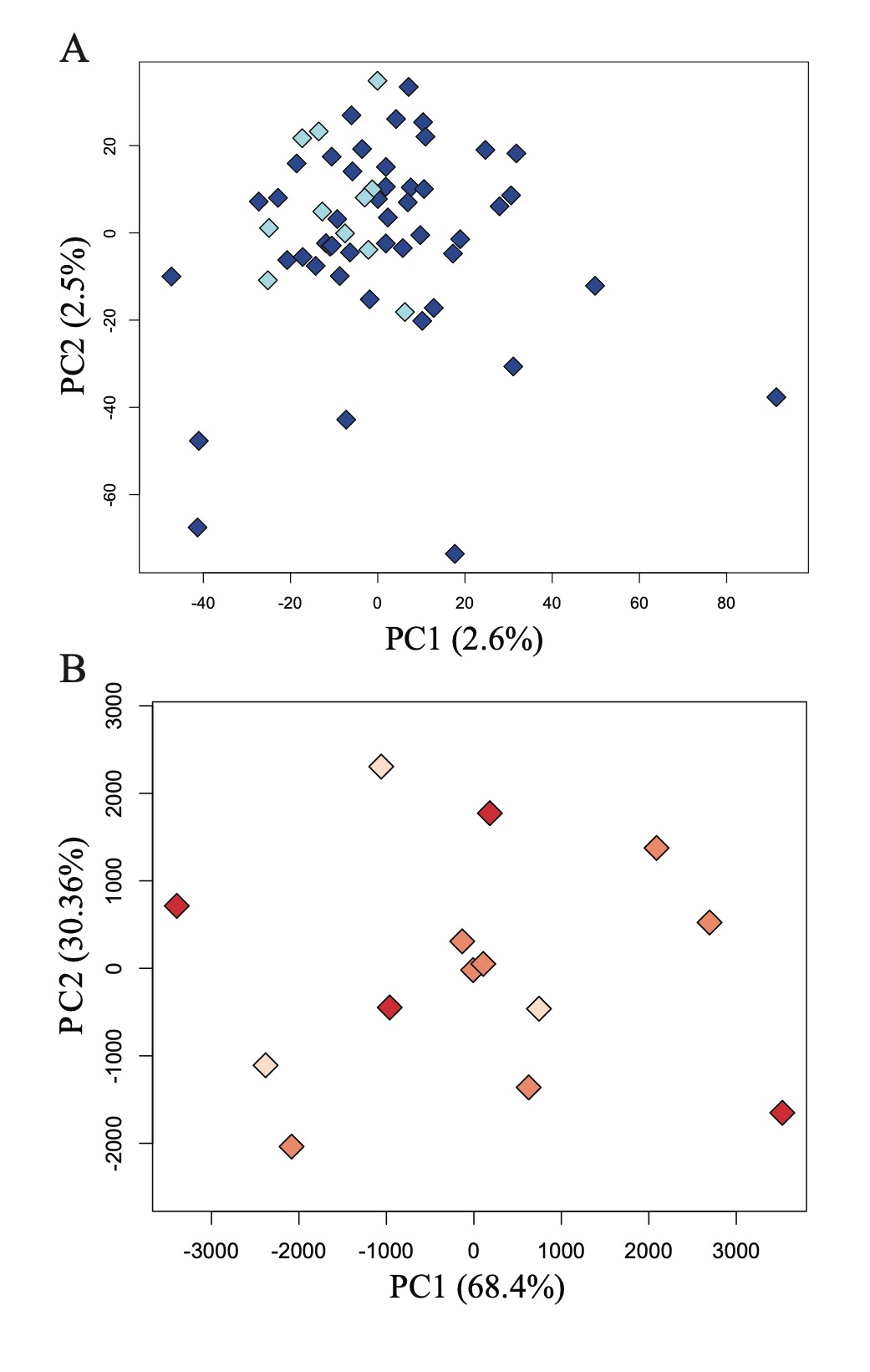

**Figure S3**. **A:** Principal Components Analysis of within-population variation among Pacific Wrens. Samples from the ranges of two subspecies, *T. p. pacificus* (dark blue) and *T. p. salebrosus* (light blue) cannot be distinguished by PCA of this data, and samples from the same locality do not cluster together. **B:** Principal Components Analysis of within-population variation among Winter Wrens. East Coast samples (dark red) are interspersed with samples from Lesser Slave Lake, Alberta (orange) and Tumbler Ridge, BC (light pink). Sex-linked loci mapping to the *Ficedula* chromosome 8 and Z chromosome have been removed to isolate patterns of geographic variation from sex-linked variation.

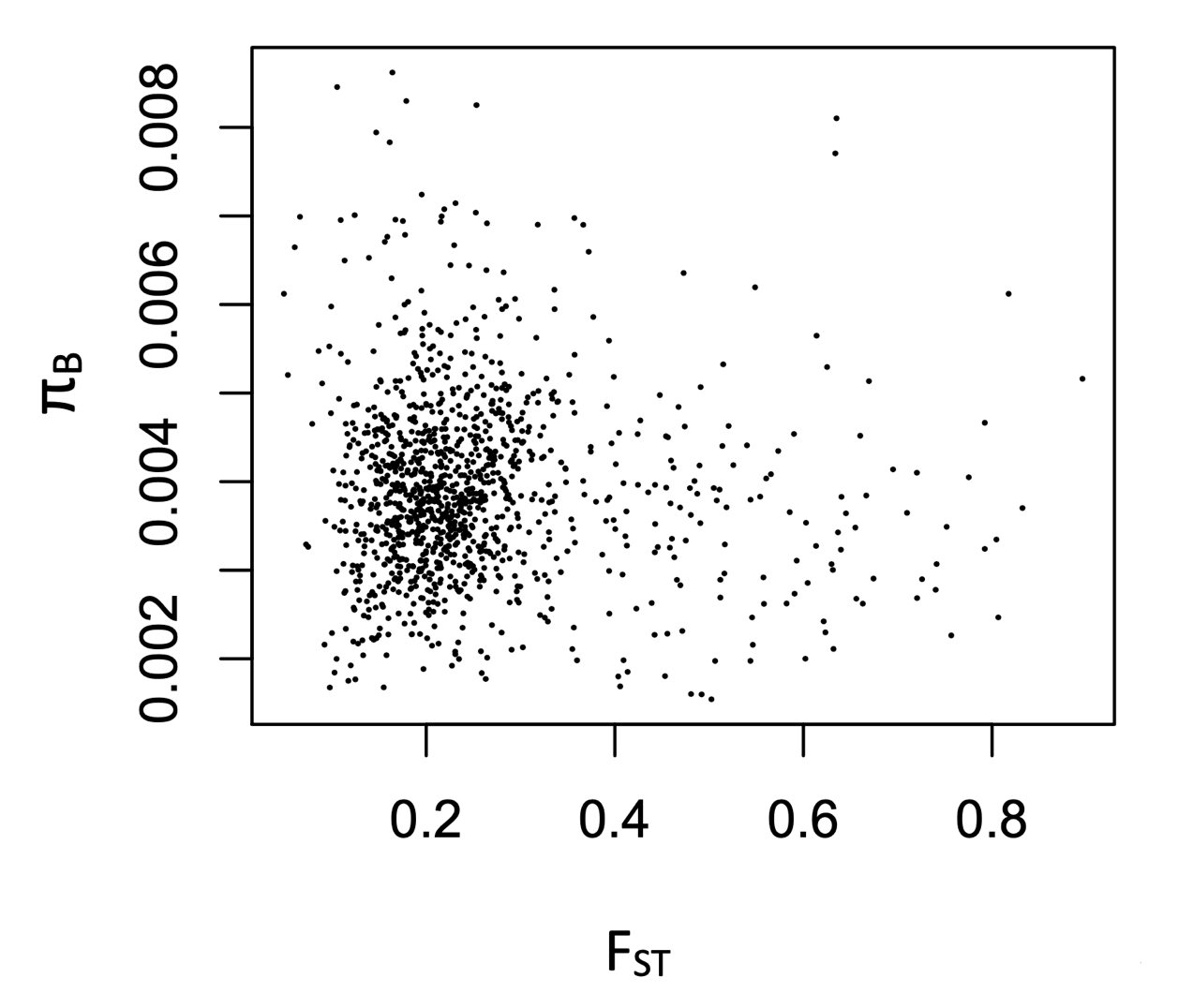

**Figure S4**. Scatterplot of *F*_ST_ vs π_B_ for 10 000-marker sliding windows between Pacific and Winter Wrens. A negative correlation is marginally significant at p=0.052.

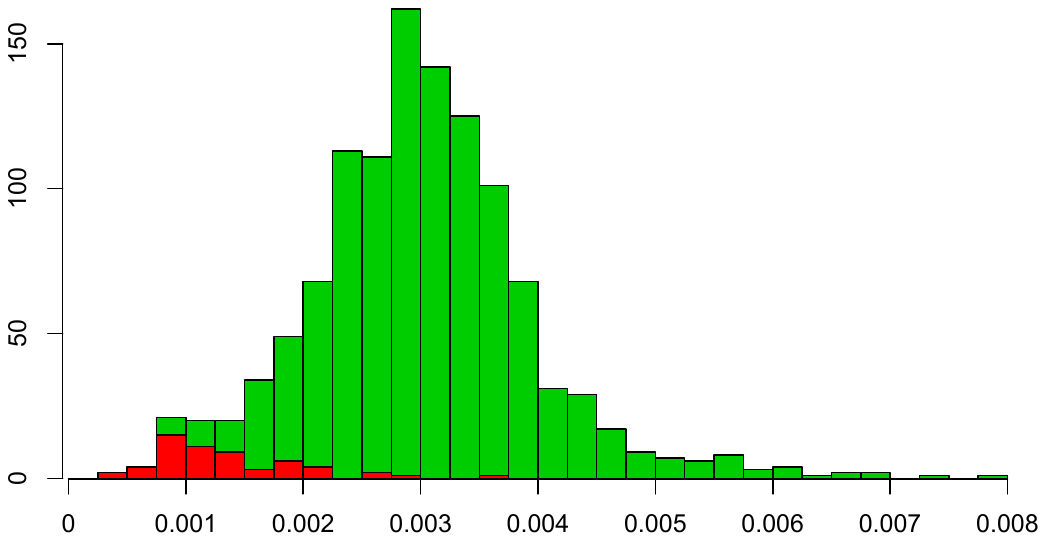

π_W_

Frequency

**Figure** **S5**. Histogram of mean π_W_ (within-species sequence diversity, averaged between the two species) for 10 000-marker sliding windows in Pacific and Winter Wrens. The 5% most-differentiated windows with the highest *F*_ST_ between Pacific and Winter Wrens are colored red.

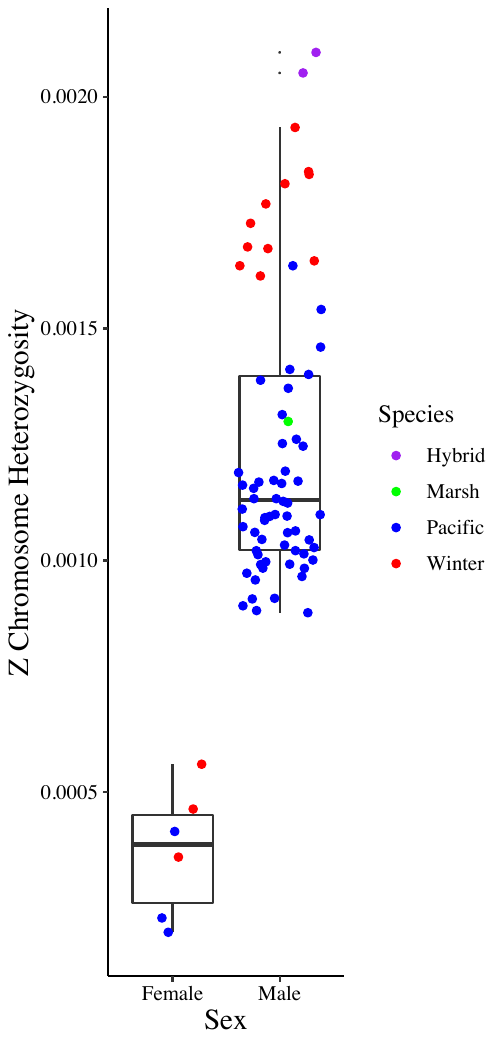

**Figure S6.** Apparent heterozygosity of the Z chromosome observed in female and male wren samples. Heterozygosity was calculated as the proportion of non-missing genotypes that were called as heterozygous in each sample at both variant and invariant sites across chromosome Z (422,161 sites in total). Females carry only one copy of the Z chromosome sequence (except for within the pseudoautosomal region) and show a greatly reduced apparent heterozygosity on chromosome Z relative to males.

**Table S4.** Synteny of gene order in the region of chromosome 8 that is inferred to have been duplicated onto the W chromosome in the wren lineage. Each column lists the genes that have been annotated in another bird species the region homologous to the inferred duplicated region. The region inferred to be duplicated in the wrens is highlighted in bold italic font. Gene synteny within the duplicated region and the flanking region is broadly conserved in all six taxa, suggesting that the arrangement on *Ficedula* chromosome 8 is the ancestral location of the duplication observed in the wrens.

| *Corvus brachyrhynchos* | *Gallus gallus* | *Taeniopygia guttata* | *Ficedula albicollis* | *Lepidothrix coronata* | *Empidonax trailli* |
| --- | --- | --- | --- | --- | --- |
| GCF_000691975.1 | GCF_000002315.5 | GCF_008822105.2 | GCF_000247815.1 | GCF_001604755.1 | GCF_003031625.1 |
|  | MIR1761 | ADGRL2 | ADGRL2 | ZRANB2 |  |
| TTLL7 | TTLL7 | TTLL7 | TTLL7 | NEGR1 |  |
| PRKACB | PRKACB | PRKACB | PRKACB | FPGT |  |
| SAMD13 | SAMD13 | SAMD13 | SAMD13 | ERICH3 |  |
| DNASE2B | DNASE2B | DNASE2B | DNASE2B | CRYZ |  |
| RPF1 | RPF1 | RPF1 | RPF1 | TYW3 |  |
| GNG5 | GNG5 | GNG5 |  | LHX8 |  |
| SPATA1 | SPATA1 | SPATA1 | SPATA1 | SLC44A5 |  |
| CTBS | CTBS | CTBS | CTBS | ACADM |  |
|  | VTG3 |  |  | RABGGTB |  |
|  | VTG2 | VIT2 |  | MSH4 |  |
|  | MIR1589 |  |  | NAA20 |  |
| SSX2IP | SSX2IP | SSX2IP | SSX2IP |  | SCAFFOLD BREAKPOINT |
| LPAR3 | LPAR3 | LPAR3 | LPAR3 | LPAR3 | LPAR3 |
| MCOLN2 | MCOLN2 | MCOLN2 | MCOLN2 | MCOLN2 | MCOLN2 |
| MCOLN3 | WDR63 | MCOLN3 | MCOLN3 |  | MCOLN3 |
| WDR63 | MCOLN3 | WDR63 | WDR63 | WDR63 | WDR63 |
| SYDE2 | SYDE2 | SYDE2 | SYDE2 | SYDE2 | SYDE2 |
|  | C8H1orf52 | C8H1orf52 | C8H1orf52 | CUNH1orf52 | CUNH1orf52 |
| BCL10 | BCL10 | BCL10 | BCL10 | BCL10 | BCL10 |
| DDAH1 | DDAH1 | DDAH1 | DDAH1 | DDAH1 | DDAH1 |
| CYR61 | CYR61 | CCN1 | CYR61 | CYR61 | CCN1 |
| ZNHIT6 | ZNHIT6 | ZNHIT6 | ZNHIT6 | ZNHIT6 | ZNHIT6 |
| COL24A1 | COL24A1 | COL24A1 | COL24A1 | COL24A1 | COL24A1 |
| ODF2L | ODF2L | ODF2L | ODF2L | ODF2L | ODF2L |
|  | CLCA2 |  |  |  |  |
|  | CLCA1 |  |  |  |  |
| SH3GLB1 | SH3GLB1 | SH3GLB1 | SH3GLB1 | SH3GLB1 | SH3GLB1 |
|  |  |  |  |  | HS2ST1 |
|  | SELENOF | SELENOF |  |  | SELENOF |
| ***HS2ST1*** | ***HS2ST1*** | ***HS2ST1*** | ***HS2ST1*** | ***HS2ST1*** | ***HS2ST1*** |
| ***LMO4*** | ***LMO4*** | ***LMO4*** | ***LMO4*** | ***LMO4*** | ***LMO4*** |
|  |  |  |  |  | ***tgfbr*** |
| ***PKN2*** | ***PKN2*** | ***PKN2*** | ***PKN2*** | ***PKN2*** | ***PKN2*** |
| ***GTF2B*** | ***GTF2B*** | ***GTF2B*** | ***GTF2B*** | ***GTF2B*** | ***GTF2B*** |
| ***KYAT3*** | ***KYAT3*** | ***KYAT3*** | ***CCBL2*** | ***KYAT3*** | ***KYAT3*** |
| ***LRRC8B*** |  | ***LRRC8B*** | ***LRRC8B*** | ***LRRC8B*** | ***LRRC8B*** |
| ***LRRC8C*** | ***LRRC8C*** | ***LRRC8C*** | ***LRRC8C*** | ***LRRC8C*** | ***LRRC8C*** |
| ***LRRC8D*** | ***LRRC8D*** | ***LRRC8D*** | ***LRRC8D*** | ***LRRC8D*** | ***LRRC8D*** |
| ***ZNF326*** | ***ZNF326*** | ***ZNF326*** | ***ZNF326*** | ***ZNF326*** | ***ZNF326*** |
|  |  |  |  |  | ***SCAFFOLD BREAKPOINT*** |
| ***BARHL2*** | ***BARHL2*** | ***BARHL2*** | ***BARHL2*** | ***BARHL2*** | ***BARHL2*** |
| ***ZNF644*** | ***ZNF644*** | ***ZNF644*** | ***ZNF644*** | ***ZNF644*** | ***ZNF644*** |
| ***HFM1*** | ***HFM1*** | ***HFM1*** | ***HFM1*** | ***HFM1*** | ***HFM1*** |
| ***CDC7*** | ***CDC7*** | ***CDC7*** | ***CDC7*** | ***CDC7*** | ***CDC7*** |
|  |  |  |  | ***SCAFFOLD BREAKPOINT*** | |
| ***TGFBR3*** | ***TGFBR3*** | ***TGFBR3*** | ***TGFBR3*** | ***TGFBR3*** | ***TGFBR3*** |
| ***BRDT*** | ***BRDT*** | ***BRDT*** | ***BRDT*** | ***BRDT*** | ***BRDT*** |
| ***EPHX4*** | ***EPHX4*** | ***EPHX4*** | ***EPHX4*** | ***EPHX4*** | ***EPHX4*** |
|  | ***KIAA1107*** | ***BTBD8*** | ***KIAA1107*** | ***KIAA1107*** | ***KIAA1107*** |
| ***CUNH1orf146*** | ***C8H1ORF146*** | ***C8H1orf146*** | ***C8H1orf146*** | ***CUNH1orf146*** | ***CUNH1orf146*** |
| ***GLMN*** | ***GLMN*** | ***GLMN*** | ***GLMN*** | ***GLMN*** | ***GLMN*** |
| ***RPAP2*** | ***RPAP2*** | ***RPAP2*** | ***RPAP2*** | ***RPAP2*** | ***RPAP2*** |
|  |  |  |  |  | ***SCAFFOLD BREAKPOINT*** |
| ***GFI1*** | ***GFI1*** | ***GFI1*** | ***GFI1*** | ***GFI1*** | ***GFI1*** |
| ***EVI5*** | ***EVI5*** | ***EVI5*** | ***EVI5*** | ***EVI5*** | ***EVI5*** |
| ***RPL5*** | ***RPL5*** | ***RPL5*** | ***RPL5*** | ***RPL5*** | ***RPL5*** |
| ***FAM69A*** | ***FAM69A*** | ***DIPK1A*** | ***FAM69A*** | ***FAM69A*** | ***DIPK1A*** |
| ***MTF2*** | ***MTF2*** | ***MTF2*** | ***MTF2*** | ***MTF2*** | ***MTF2*** |
| ***TMED5*** | ***TMED5*** | ***TMED5*** | ***TMED5*** | ***TMED5*** | ***TMED5*** |
| ***CCDC18*** | ***CCDC18*** | ***CCDC18*** | ***CCDC18*** | ***CCDC18*** | ***CCDC18*** |
|  | ***RDH8*** |  |  |  |  |
| ***DR1*** | ***DR1*** | ***DR1*** | ***DR1*** | ***DR1*** | ***DR1*** |
| ***FNBP1L*** | ***FNBP1L*** | ***FNBP1L*** | ***FNBP1L*** | ***FNBP1L*** |  |
|  | ***TRNAR-UCU*** | |  |  |  |
| ***BCAR3*** | ***BCAR3*** | ***BCAR3*** | ***BCAR3*** | ***BCAR3*** | ***BCAR3*** |
| ***TRNAR-UCU*** | | ***TRNAR-UCU*** | ***TRNAR-UCU*** | ***TRNAR-UCU*** | ***TRNAR-UCU*** |
| ***DNTTIP2*** | ***DNTTIP2*** | ***DNTTIP2*** | ***DNTTIP2*** | ***DNTTIP2*** | ***DNTTIP2*** |
| ***GCLM*** | ***GCLM*** | ***GCLM*** | ***GCLM*** | ***GCLM*** | ***GCLM*** |
| ***TECR*** | ***TECR*** |  | ***TECR*** | ***TECR*** | ***TECR*** |
| ***ABCA4*** | ***ABCA4*** | ***ABCA4*** | ***ABCA4*** | ***ABCA4*** | ***ABCA4*** |
| ***ARHGAP29*** | ***ARHGAP29*** | ***ARHGAP29*** | ***ARHGAP29*** | ***ARHGAP29*** | ***ARHGAP29*** |
| ***ABCD3*** | ***ABCD3*** | ***ABCD3*** | ***ABCD3*** | ***ABCD3*** | ***ABCD3*** |
| ***F3*** | ***F3*** | ***F3*** | ***SLC44A3*** | ***F3*** | ***F3*** |
|  |  |  |  | ***SCAFFOLD BREAKPOINT*** | |
| ***SLC44A3*** | ***SLC44A3*** | ***SLC44A3*** | ***F3*** | ***SLC44A3*** | ***SLC44A3*** |
| ***CNN3*** | ***CNN3*** | ***CNN3*** | ***CNN3*** | ***CNN3*** | ***CNN3*** |
| ***ALG14*** | ***ALG14*** | ***ALG14*** | ***ALG14*** | ***ALG14*** | ***ALG14*** |
| ***TMEM56*** | ***TMEM56*** |  | ***TMEM56*** |  | ***TMEM56*** |
|  | ***HCCS*** | ***TLCD4*** |  |  |  |
| ***RWDD3*** | ***RWDD3*** | ***RWDD3*** | ***RWDD3*** | ***RWDD3*** | ***RWDD3*** |
| ***PTBP2*** | ***PTBP2*** | ***PTBP2*** | ***PTBP2*** | ***PTBP2*** | ***PTBP2*** |
| ***DPYD*** | ***DPYD*** | ***DPYD*** | ***DPYD*** | ***DPYD*** | ***DPYD*** |
|  | ***MIR137*** | ***MIR137*** |  |  |  |
| ***SNX7*** | ***SNX7*** | ***SNX7*** | ***SNX7*** | ***SNX7*** | ***SNX7*** |
| ***PLPPR5*** | ***PLPPR4*** | ***PLPPR5*** | ***PLPPR5*** | ***PLPPR5*** | ***PLPPR5*** |
| ***PLPPR4*** | ***PLPPR5*** | ***PLPPR4*** | ***PLPPR4*** | ***PLPPR4*** | ***PLPPR4*** |
| ***PALMD*** | ***PALMD*** | ***PALMD*** | ***PALMD*** | ***PALMD*** | ***PALMD*** |
| ***FRRS1*** | ***FRRS1*** | ***FRRS1*** | ***FRRS1*** | ***FRRS1*** | ***SCAFFOLD BREAKPOINT*** |
| ***AGL*** | ***AGL*** | ***AGL*** | ***AGL*** | ***AGL*** | ***AGL*** |
| ***SLC35A3*** | ***SLC35A3*** | ***SLC35A3*** | ***SLC35A3*** | ***SLC35A3*** | ***SLC35A3*** |
| ***MFSD14A*** | ***MFSD14A*** | ***MFSD14A*** | ***MFSD14A*** | ***MFSD14A*** | ***MFSD14A*** |
| ***SASS6*** | ***SASS6*** | ***SASS6*** | ***SASS6*** | ***SASS6*** | ***SASS6*** |
| ***TRMT13*** | ***TRMT13*** | ***TRMT13*** | ***TRMT13*** | ***TRMT13*** | ***TRMT13*** |
| ***LRRC39*** | ***LRRC39*** | ***LRRC39*** | ***LRRC39*** | ***LRRC39*** | ***LRRC39*** |
| ***DBT*** | ***DBT*** | ***DBT*** | ***DBT*** | ***DBT*** | ***DBT*** |
| ***RTCA*** | ***RTCA*** | ***RTCA*** | ***RTCA*** | ***RTCA*** | ***RTCA*** |
| ***VCAM1*** | ***VCAM1*** | ***VCAM1*** | ***VCAM1*** | ***VCAM1*** | ***VCAM1*** |
| ***GPR88*** |  | ***GPR88*** | ***GPR88*** | ***GPR88*** | ***GPR88*** |
| ***CDC14A*** | ***CDC14A*** | ***CDC14A*** |  | ***CDC14A*** | ***CDC14A*** |
| ***EXTL2*** | ***EXTL2*** | ***EXTL2*** | ***EXTL2*** | ***EXTL2*** | ***EXTL2*** |
| ***SLC30A7*** | ***SLC30A7*** | ***SLC30A7*** | ***SLC30A7*** | ***SLC30A7*** | ***SLC30A7*** |
| ***DPH5*** | ***DPH5*** | ***DPH5*** | ***DPH5*** | ***DPH5*** | ***DPH5*** |
|  | ***MIR1610*** |  |  |  |  |
| ***S1PR1*** | ***S1PR1*** | ***S1PR1*** | ***S1PR1*** | ***S1PR1*** | ***S1PR1*** |
| ***OLFM3*** | ***OLFM3*** | ***OLFM3*** | ***OLFM3*** | ***OLFM3*** | ***OLFM3*** |
| COL11A1 | COL11A1 | COL11A1 | COL11A1 | COL11A1 | COL11A1 |
| RNPC3 | RNPC3 | RNPC3 | RNPC3 | RNPC3 | RNPC3 |
|  |  |  |  | SCAFFOLD BREAKPOINT | SCAFFOLD BREAKPOINT |
| NTNG1 | NTNG1 | NTNG1 |  | NTNG1 | NTNG1 |
| VAV3 | VAV3 | VAV3 |  | VAV3 | VAV3 |
| SLC25A24 | SLC25A24 | SLC25A24 | SLC25A24 | SLC25A24 | SLC25A24 |
| FAM102B | FAM102B | FAM102B | FAM102B | FAM102B | FAM102B |
| HENMT1 | HENMT1 | HENMT1 | HENMT1 | HENMT1 | HENMT1 |
|  |  |  |  | SCAFFOLD BREAKPOINT | |
| PRPF38B | PRPF38B | PRPF38B | PRPF38B | PRPF38B | PRPF38B |
| FNDC7 | FNDC7 | FNDC7 | FNDC7 | FNDC7 | FNDC7 |
| STXBP3 | STXBP3 | STXBP3 | STXBP3 | STXBP3 | STXBP3 |
| AKNAD1 | GPSM2 | AKNAD1 | AKNAD1 | AKNAD1 | AKNAD1 |
| GPSM2 | AKNAD1 | GPSM2 | GPSM2 | GPSM2 | GPSM2 |
| CLCC1 | CLCC1 | CLCC1 | CLCC1 | CLCC1 | CLCC1 |
| WDR47 | WDR47 | WDR47 | WDR47 | WDR47 | WDR47 |
| CAMSAP2 | CAMSAP2 | CAMSAP2 | CAMSAP2 | CAMSAP2 | CAMSAP2 |
| DDX59 | DDX59 | DDX59 | DDX59 | DDX59 | DDX59 |
| KIF14 | KIF14 | KIF14 | KIF14 | KIF14 | KIF14 |

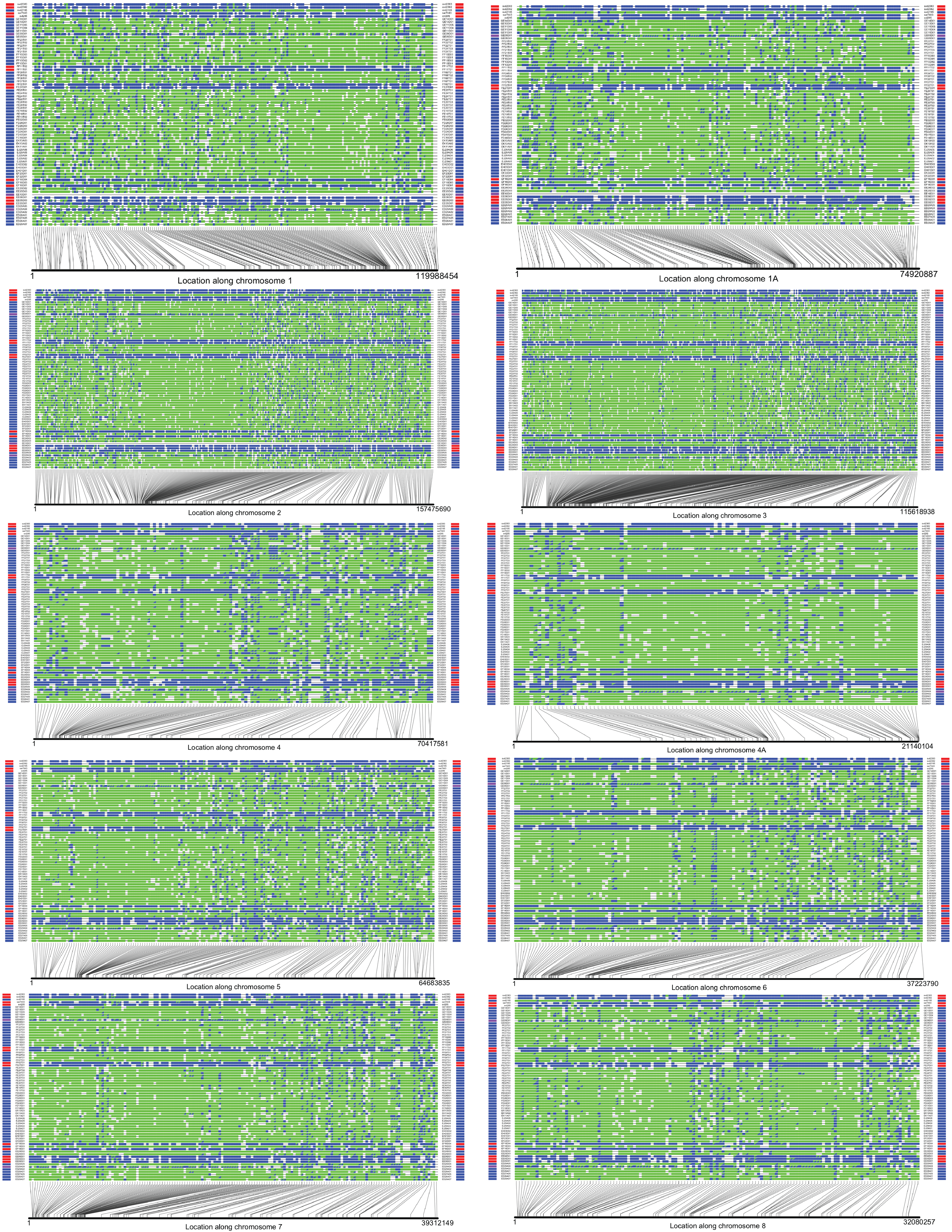

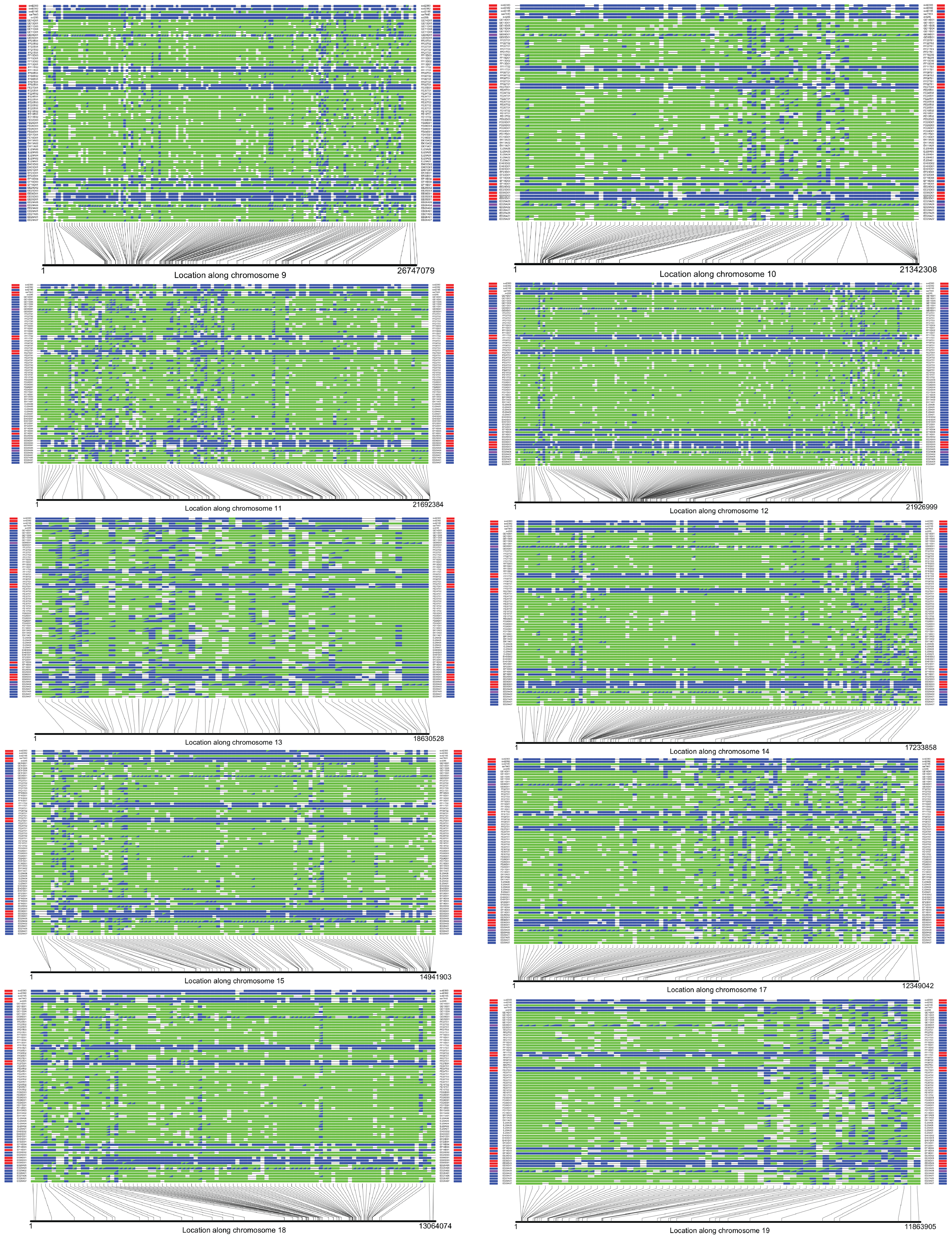

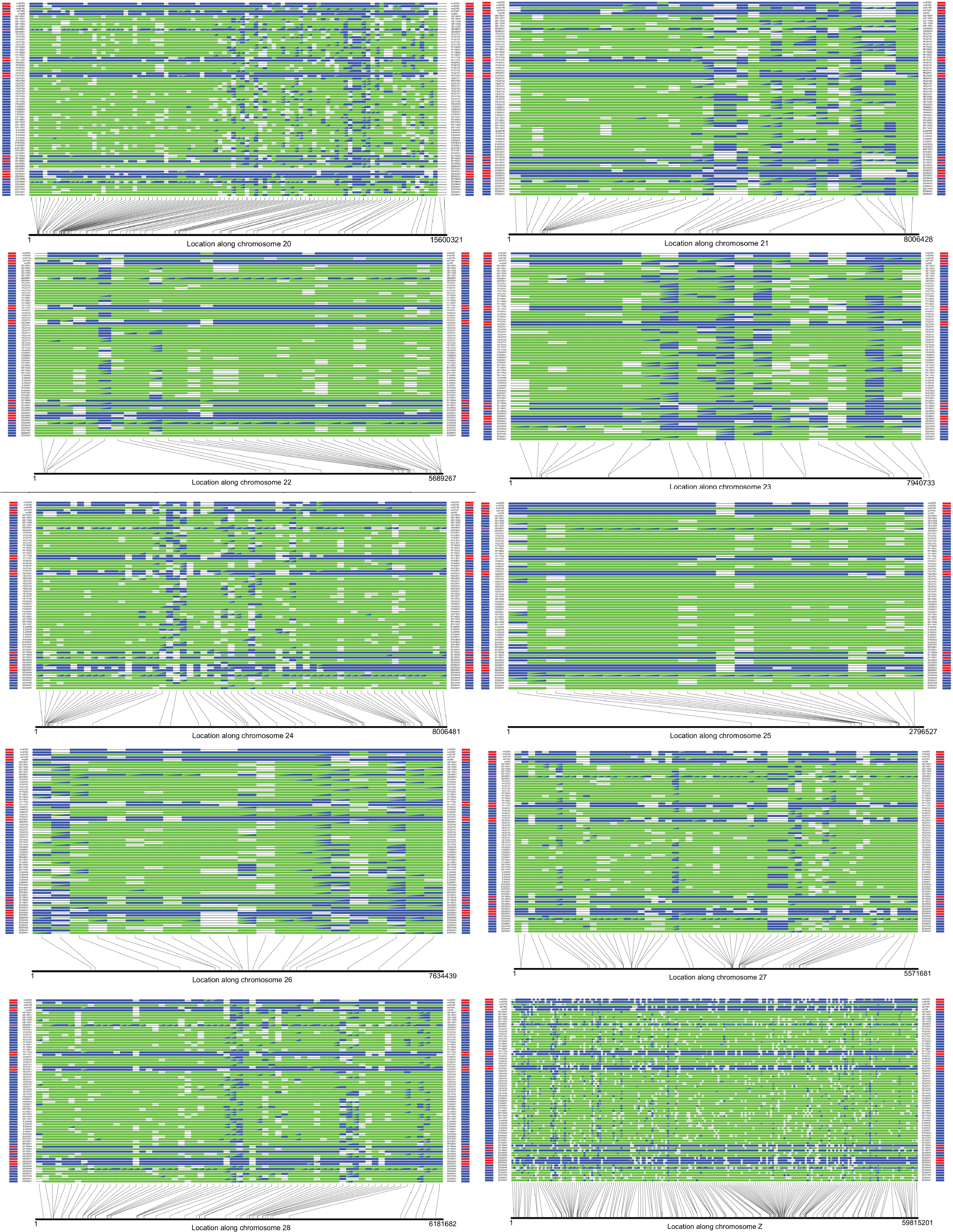

**Figure S7.** Scan for introgressed loci. Genotypes for each sample are plotted in rows, with sample names labelled at the start and end of each row. Colours adjacent to the sample names indicate species identity: Winter Wren (red), Pacific Wren (blue), hybrid (purple), or Marsh Wren (grey). Each column represents a SNP with *F*_ST_ greater than 0.9. The genotype of each sample at each locus is indicated by a green box (homozygous for the Pacific Wren-associated allele), blue box (Winter Wren-associated allele), or blue and green triangles (heterozygous). Winter Wrens samples appear as blue stripes on the plots, while the two hybrids are heterozygous at many loci. A short introgressed block is visible on the 5’ (left) end of chromosome 24 in Winter Wren sample EF18D01, where it contains a run of Pacific Wren-associated alleles. Full resolution images can be viewed at the Github and Dryad repositories.
